# Exploring the multifactorial causes and therapeutic strategies for anabolic resistance in sarcopenia: A systems modeling study

**DOI:** 10.1101/2025.09.12.675977

**Authors:** Taylor J. McColl, Daniel R. Moore, Eldon Emberly, David D. Church, David C. Clarke

## Abstract

**Background:** Sarcopenia is the progressive loss of skeletal muscle mass, strength, and function with age, driven by dysregulation in the rates of muscle protein synthesis (MPS) and breakdown (MPB). Although MPB contributes to net protein balance (NB), a primary contributor of sarcopenia is *anabolic resistance*, defined as the blunted MPS response to anabolic stimuli such as feeding. While candidate mechanisms of anabolic resistance have been identified, none singularly accounts for the observed reduction in MPS. Instead, multiple mechanisms likely act simultaneously and interactively to suppress MPS. Studying these interactions experimentally is challenging. Mathematical modeling is well suited to analyzing complex biological phenomena such as anabolic resistance.

**Methods:** We analyzed a previously developed kinetic model of leucine-mediated signaling and protein metabolism in human skeletal muscle to systematically investigate potential mechanisms contributing to anabolic resistance. Using global sensitivity analysis, we identified key controllers of MPS, MPB, and NB. We then simulated amino acid feeding in older adults, classified the responses as either anabolic sensitive or resistant, and compared the resulting parameter distributions of the two groups. We next performed targeted analysis to evaluate the effects of individual and combined putative mechanisms of anabolic resistance on muscle metabolism. Finally, we simulated therapeutic interventions aimed at restoring muscle metabolism.

**Results:** The sensitivity analysis revealed that MPS and MPB are primarily controlled by their proximal signaling processes, while NB is largely driven by MPS dynamics. Exploratory simulations showed that several parameters and signaling protein concentrations, particularly those controlling MPS, differed significantly between anabolic sensitive and resistant groups. The targeted simulations indicated that multiple dysregulated mechanisms were required to account for the experimentally observed reductions in MPS in older adults. Therapy simulations showed that single-target interventions could largely restore MPS when isolated mechanisms were perturbed (e.g., increasing mTORC1 sensitivity, enhancing p70S6K levels), but a multifactorial approach was required to recover muscle metabolism when all anabolic resistance mechanisms were present.

**Conclusion:** This study highlights the multifactorial nature of anabolic resistance and the implications for therapy. Specifically, the results motivate new hypotheses regarding the mechanisms most likely to be impaired and argue for multi-target therapeutic strategies to help restore muscle protein metabolism in aging.

## Introduction

Sarcopenia is the progressive decline in skeletal muscle mass, strength, and function that accompanies aging [1]. Those with sarcopenia experience higher hospitalization rates [2] and contribute to increased public healthcare costs [3–5]. Given the impact of sarcopenia on health and quality of life, further research is needed to enhance our understanding of age-related muscle loss and to develop effective strategies for its prevention and management.

Skeletal muscle mass is largely determined by its protein content, which undergoes continuous turnover through concurrent synthesis and degradation processes [6]. The net balance (NB) between the rates of muscle protein synthesis (MPS) and muscle protein breakdown (MPB) governs overall muscle protein levels. MPS increases in response to anabolic stimuli such as nutrient intake [7], with amino acid availability – particularly leucine [8–10] – playing a key stimulatory role. In contrast, the attenuation of MPB during the postprandial period is primarily mediated via an increase in plasma insulin levels [11]. In the postabsorptive state, MPS and MPB rates are generally similar between young and older adults [12–15]. However, aging muscle exhibits *anabolic resistance*, characterized by a blunted stimulation of MPS and impaired suppression of MPB following anabolic stimuli [16–18]. Over time, this anabolic resistance is thought to be the primary driver of the gradual loss of skeletal muscle with age.

Anabolic resistance of aging is thought to be caused by a range of factors including reduced amino acid availability to muscle due to increased splanchnic extraction [19–21], impaired transport from blood into muscle [20,22], impaired insulin-mediated protein signaling [23,24], reduced mTOR and p70S6K protein levels [18,25], and diminished mTORC1 signaling sensitivity to protein intake [18,26]. Sustained MPB rates are due to reduced insulin-mediated blunting of proteolysis or other age-related factors, such as chronic inflammation [27–29]. Effectively understanding and treating anabolic resistance therefore requires quantifying the relative contributions of these mechanisms and identifying targets for intervention.

The limited long-term success of treatments for anabolic resistance may stem from the simultaneous dysregulation of multiple mechanisms affecting muscle protein metabolism. Nutritional and pharmacological interventions tend to target individual mechanisms or fail to target the causal mechanisms altogether. Therefore, therapeutic strategies that target the multiple interacting causal mechanisms may be needed.

Mathematical models are valuable tools for studying complex biological systems. They could be applied to identify the most likely combination of dysregulated mechanisms underlying anabolic resistance in older adults to inform the human clinical research needed to fully characterize the myriad potential biological mediators. We recently developed a kinetic model of leucine-mediated skeletal muscle signaling and protein metabolism in human skeletal muscle, incorporating key leucine- and insulin-mediated pathways that control muscle protein metabolism [10]. Although originally developed using data from healthy younger adults, these molecular pathways are also present in older individuals but likely operate with altered reaction rates. Therefore, this kinetic model provides a framework to investigate the multifactorial nature of anabolic resistance and characterize how pathophysiological mechanisms drive impaired feeding-induced muscle metabolism.

The purpose of this study was to investigate the multifactorial causes of anabolic resistance by quantifying how dysregulated mechanisms in muscle protein metabolism contribute to impaired muscle mass in older adults. Specifically, we applied sensitivity analyses to the McColl and Clarke kinetic model [10] to identify key parameters controlling muscle metabolism. We pursued two strategies: 1) a “naïve” global sensitivity analysis to identify parameters potentially responsible for reduced MPS, and 2) a targeted sensitivity analysis that simulated previously proposed anabolic resistance mechanisms. These targeted simulations were then used to evaluate therapeutic strategies aimed at restoring MPS and NB. Collectively, our findings provide insight into the principal physiological drivers of anabolic resistance and identify promising targets for therapeutic intervention to mitigate muscle loss in older adults.

## Methods

### Kinetic model of leucine-mediated signaling and muscle protein metabolism

The McColl and Clarke kinetic model of leucine-mediated signaling and muscle protein metabolism in human skeletal muscle [10] (henceforth, the kinetic model) served as the foundation for this study (Figure 1). This model simulates skeletal muscle protein metabolism in response to leucine ingestion in healthy adults and contains four interconnected modules: 1) the mTOR signaling module, 2) the leucine kinetic module, 3) the digestive system module, and 4) the insulin secretion module [10]. Together, these modules simulate the ingestion of leucine, leucine-mediated stimulation of insulin-and leucine-dependent signaling pathways, and muscle protein metabolism. While originally developed using data from healthy adults, the model can be applied to study anabolic resistance in sarcopenia because the biological mechanisms it represents are conserved in young and older individuals. Differences between young and older adults manifest as differences in parameter values. Further details of the kinetic model are provided in the Supplementary Information.

**Figure 1:**
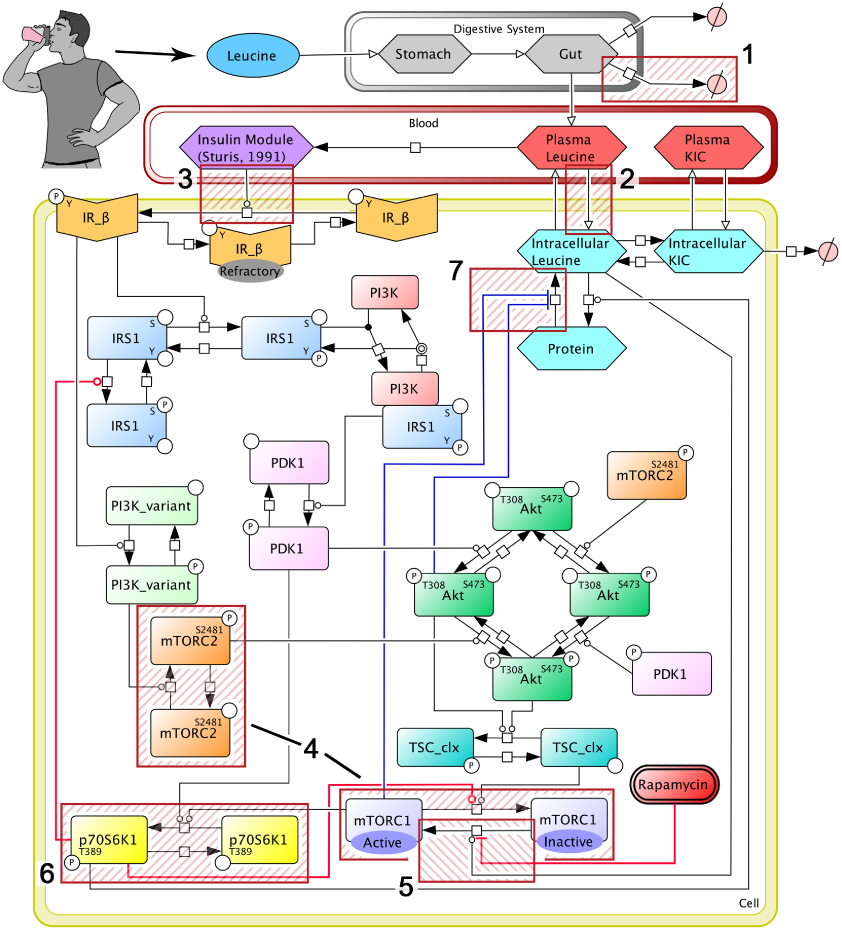
Kinetic model of leucine-mediated signalling and muscle protein metabolism. The topology of the McColl and Clarke kinetic model is shown, with hatched red boxes highlighting the investigated mechanisms of anabolic resistance: 1) increased first-pass splanchnic extraction, 2) reduced transport of amino acids into muscle, 3) reduced insulin-mediated signalling due to insulin resistance, 4) reduced mTOR concentration, 5) reduced sensitivity of mTORC1 to leucine signaling, 6) reduced p70S6K concentration, and 7) elevated muscle protein breakdown rates.

### Defining anabolic resistance

We established an anabolic resistance “threshold” based on two studies by Mitchell et al. [30,31]. Although these studies measured myofibrillar protein synthesis specifically, this protein fraction, which constitutes ∼60% of muscle proteins, demonstrates a similar nutrient sensitivity and age-related resistance as sarcoplasmic proteins [18,32]. Both studies used the same intervention: young [30] and older adults [31] received a 15-gram mixed essential amino acid (EAA) bolus (3.59 grams of leucine), with muscle biopsies collected over the subsequent four hours to calculate MPS, measured as fractional synthetic rate (FSR) [30,31]. Using the “total leucine synthesized” calculations from McColl and Clarke [10], the experimentally measured MPS response in young adults was 0.46±0.11 grams of leucine synthesized in muscle over four hours, while in older adults it was 0.27±0.09 grams of leucine. Despite the older adults being classified as healthy [31], the reduced MPS response to EAA feeding indicated the presence of anabolic resistance. To define the anabolic resistance threshold, we used the lower 95% confidence interval of the younger adult MPS response (0.46±0.11 grams of leucine), yielding a threshold of 0.25 g leucine synthesized in muscle. Simulated MPS responses below this threshold were classified as *anabolically resistant*, whereas those above it were considered *anabolically sensitive*. This value closely approximates the mean MPS response for the older adult group.

### Multi-parameter sensitivity analysis

We applied a multi-parameter sensitivity analysis (MPSA) to generate physiologically plausible parameter sets, with the purpose of identifying model parameters that most influence system-level behaviors representative of anabolic resistance. MPSA is a global sensitivity analysis approach that systematically samples parameter sets and evaluates their relative contribution to model dynamics [33]. Complete details for the MPSA, including the criteria for parameter filtering and analyses applied to the filtered parameter sets, are provided in the Supplementary Information.

#### MPSA: Kolmogorov-Smirnov test

We applied the two-sample Kolmogorov-Smirnov (KS) test to assess whether parameter sets from the anabolic-sensitive and anabolic-resistant groups originated from the same distribution or represented unique distributions. We performed the test using the R function *ks.test*, which calculated the KS statistic and the associated p-value. To account for multiple comparisons, we applied false discovery rate (FDR) correction using the Benjamini-Hochberg method (R function: *p.adjust*) to compute the *q*-value. A *q*-value less than 0.05 indicated a statistically significant difference between parameter distributions.

### Targeted approach to evaluate the contributions of anabolic resistance mechanisms to muscle protein metabolism

We used a targeted approach to evaluate how the putative mechanisms underlying anabolic resistance in sarcopenia independently influence muscle protein metabolism and how these mechanisms might operate synergistically to further impair muscle protein metabolism. As noted in the Introduction and outlined in Table 1, several mechanisms contribute to anabolic resistance (Figure 1). Of note, elevated postabsorptive levels of phospho-p70S6K have been observed in older adults, which we hypothesized reflects a compensatory response to dysregulated upstream mechanisms [10,34] – a phenomenon we term “compensatory signaling”.

**Table 1.**
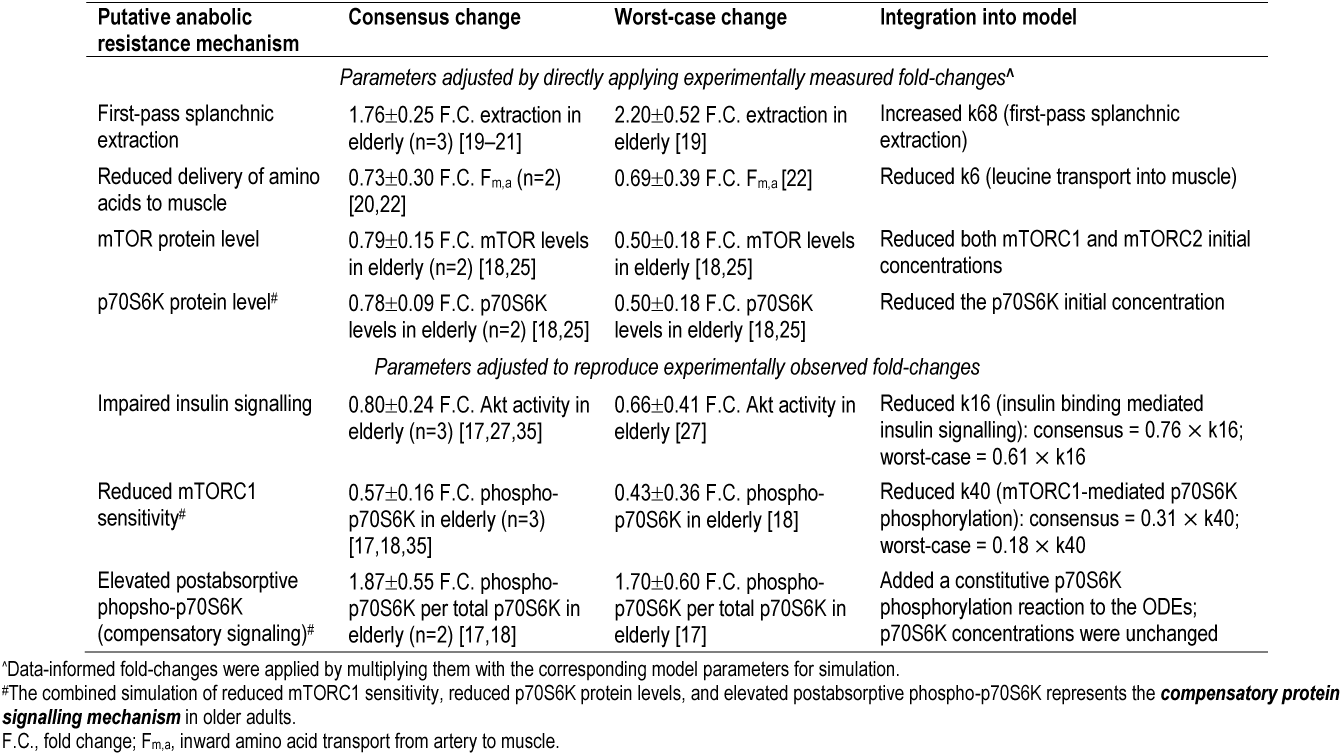
Modeling the putative anabolic resistance mechanisms. Summary of putative mechanisms contributing to anabolic resistance in sarcopenia, along with quantitative evidence for the consensus (i.e., average) and worst-case effects identified across a subset of studies. The worst-case effect was selected from the study reporting the greatest deviation in fold-change relative to the young group. The number of studies informing each consensus estimate is indicated by n. The kinetic model parameters that were altered to represent the corresponding mechanism are listed in the rightmost column.

| Putative anabolic resistance mechanism | Consensus change | Worst-case change | Integration into model |
| --- | --- | --- | --- |
| *Parameters adjusted by directly applying experimentally measured fold-changes<sup>a</sup>* |  |  |  |
| First-pass splanchnic extraction | 1.76±0.25 F.C. extraction in elderly (n=3) [19–21] | 2.20±0.52 F.C. extraction in elderly [19] | Increased k68 (first-pass splanchnic extraction) |
| Reduced delivery of amino acids to muscle | 0.73±0.30 F.C. $F_{m,a}$ (n=2) [20,22] | 0.69±0.39 F.C. $F_{m,a}$ [22] | Reduced k6 (leucine transport into muscle) |
| mTOR protein level | 0.79±0.15 F.C. mTOR levels in elderly (n=2) [18,25] | 0.50±0.18 F.C. mTOR levels in elderly [18,25] | Reduced both mTORC1 and mTORC2 initial concentrations |
| p70S6K protein level<sup>#</sup> | 0.78±0.09 F.C. p70S6K levels in elderly (n=2) [18,25] | 0.50±0.18 F.C. p70S6K levels in elderly [18,25] | Reduced the p70S6K initial concentration |
| *Parameters adjusted to reproduce experimentally observed fold-changes* |  |  |  |
| Impaired insulin signalling | 0.80±0.24 F.C. Akt activity in elderly (n=3) [17,27,35] | 0.66±0.41 F.C. Akt activity in elderly [27] | Reduced k16 (insulin binding mediated insulin signalling): consensus = $0.76 \times k16$; worst-case = $0.61 \times k16$ |
| Reduced mTORC1 sensitivity<sup>#</sup> | 0.57±0.16 F.C. phospho-p70S6K in elderly (n=3) [17,18,35] | 0.43±0.36 F.C. phospho-p70S6K in elderly [18] | Reduced k40 (mTORC1-mediated p70S6K phosphorylation): consensus = $0.31 \times k40$; worst-case = $0.18 \times k40$ |
| Elevated postabsorptive phospho-p70S6K (compensatory signaling)<sup>#</sup> | 1.87±0.55 F.C. phospho-p70S6K per total p70S6K in elderly (n=2) [17,18] | 1.70±0.60 F.C. phospho-p70S6K per total p70S6K in elderly [17] | Added a constitutive p70S6K phosphorylation reaction to the ODEs; p70S6K concentrations were unchanged |

<sup>a</sup>Data-informed fold-changes were applied by multiplying them with the corresponding model parameters for simulation.
<sup>#</sup>The combined simulation of reduced mTORC1 sensitivity, reduced p70S6K protein levels, and elevated postabsorptive phospho-p70S6K represents the **compensatory protein signalling mechanism** in older adults.
F.C., fold change; $F_{m,a}$ , inward amino acid transport from artery to muscle.

For each mechanism, we curated experimental data quantifying the differences between young and older adults, identified the model parameter(s) corresponding to each mechanism, and estimated both a *consensus fold change* and a *worst-case fold change* in the values of these parameters between young and old based on a subset of relevant studies (Table 1). Several of the studies directly measured processes represented in the model, including first-pass splanchnic extraction, reduced amino acid delivery to muscle (F_m,a_), and the protein levels of mTOR and p70S6K. However, other studies characterized related signaling components rather than the specific modelled processes. In these cases, we adjusted model parameter values to reproduce the experimentally observed fold changes, ensuring that the model captured the functional consequences of these upstream alterations. This strategy was applied to impaired insulin signaling, reduced mTORC1 sensitivity, and elevated post-absorptive phospho-p70S6K (compensatory signaling).

The consensus fold change for each mechanism was calculated in two steps. First, the fold change within each study was calculated as the ratio between the mean response of the young and older adults, with the corresponding standard error (SE) derived using propagation of error. Second, a consensus mean fold change for each mechanism was calculated across studies using an inverse variance-weighted average, which assigns greater weight to estimates with higher precision (Equations 7-9) [36]:

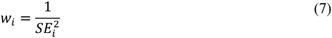

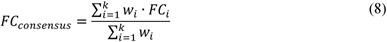

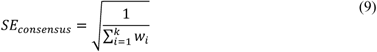

where *i* is each of the *k* independent studies, *FC_i_* and *SE_i_* are the fold change (FC) and corresponding SE for study *i*, respectively, *w*_*i*_ is the inverse variance weight of study *i*, and *FC_consensus_* and *SE_consensus_* are the consensus fold change and SE across *k* studies for the respective putative mechanism of anabolic resistance. The worst-case fold change was assigned as the value from the study within the group of studies that reported the greatest deviation from the young adult population.

We then used the kinetic model to simulate the effects of each putative mechanism individually, as well as the combined effect of all mechanisms together, in both the presence and absence of elevated postabsorptive phospho-p70S6K (i.e., compensatory signaling), as follows. First, we created a virtual population of 100 anabolic sensitive (i.e., “healthy”) models by randomly sampling 100 parameter sets from the anabolic sensitive group of the MPSA (Naïve approach). Performing the simulations for a virtual population was done to foster the generalizability of the results. Second, the anabolic resistance mechanisms were simulated within each of the 100 models by multiplying the relevant model kinetic parameter or initial protein concentration by the fold change corresponding consensus or worst-case estimate. For the simulated compensatory phospho-p70S6K signaling mechanism, an additional reaction was added to the model to simulate the elevated postabsorptive phosphorylation rate, with its kinetic parameter informed by the estimates reported in Table 1. We then simulated the anabolic resistant models in response to a 3.59 g leucine bolus and extracted the MPS, MPB, and NB responses over the 4-hour postprandial period. This dose was chosen to allow comparison with the Mitchell et al. studies, which provided a 15 g EAA bolus containing 3.59 g leucine and informed the anabolic resistance thresholds.

### Simulation of therapeutic strategies to mitigate anabolic resistance

Therapeutic strategies to mitigate anabolic resistance were simulated using models developed through the Targeted Approach, which had quantified the effects of individual or combined anabolic resistance mechanisms on MPS, MPB, and NB. Each therapeutic strategy involved restoring one or more dysregulated parameters to their “healthy” value by multiplying the parameters by the inverse of its consensus or worst-case fold-change estimate. Each therapeutic intervention was applied across all anabolic resistance simulations from the Targeted Analysis – including those in which the targeted parameter was not originally perturbed. This approach attempts to mimic the scenario in which one or more druggable targets are treated when the causal mechanism itself is not druggable (cf. [37,38]). The efficacy of each therapeutic intervention was assessed against its ability to recover MPS and NB relative to the healthy response.

### Numerical methods and software

The 95% confidence interval was calculated using one-tail and the t-value corresponding to the appropriate degrees of freedom. MATLAB version R2022a was used to perform the MPSA, model simulations, and multiple regression analyses. R (version 4.3.2) was used for statistical testing and the visualization of results using the ggplot2 package [39]. WebPlotDigitizer was used to extract published data (i.e., means, SE) that were only reported in figures [40]. The code generated in this study is freely available in the GitHub repository: https://github.com/tjmccoll/MultifactorialCauseOfAnabolicResistance/.

## Results

### Multi-parameter sensitivity analysis of the kinetic model

The MPSA resulted in 2,663 physiologically plausible parameter sets for healthy adults. Regression models using log_10_-transformed outcome measures and log_10_-transformed parameter values were the most predictive for MPS and MPB, whereas using raw outcomes with log_10_-transformed parameter values were most predictive for NB. The best-performing regression models for MPS, MPB, and NB include the top 35, 45, and 35 parameters from the main-effect-only models as pairwise interaction terms, respectively (Table S2). The p-1 sensitivity analysis of MPS and MPB revealed that parameters most proximal to their respective reactions had the greatest influence on their outcomes (Table S3). For NB, the most influential parameters were a combination of those affecting MPS and MPB, but parameters more proximal to MPS had a greater influence on the NB response (Table S3).

### Naïve approach reveals mechanisms contributing to anabolic resistance

We used an unbiased, exploratory approach to assess mechanisms that may contribute to anabolic resistance in sarcopenia and influence muscle protein metabolism. To do so, we repeated the MPSA but with the following modifications. First, we excluded the MPS criterion because the original MPSA was constrained to generate physiologically reasonable MPS responses in healthy, young adults. This criterion likely restricted the range of MPS responses observed in older adults with anabolic resistance (see “Defining Anabolic Resistance” in Methods), thereby removing relevant parameter sets. To better simulate theoretical MPS responses in older adults, we removed the constraint to allow the full breadth of MPS variability to be captured. Second, the original MPSA featured fixed values for the parameters controlling the model’s input function. However, one of these parameters controls the rate of splanchnic extraction (k_68_), which literature suggests is elevated in older adults [19–21]. To comprehensively assess mechanisms contributing to anabolic resistance, we allowed k_68_ to vary as part of the MPSA. We first performed an MPSA in which only k_68_ was varied within a two-fold range to determine its physiologically plausible range (MPS criterion removed). This range was then used to constrain k_68_ in the subsequent MPSA to ensure sufficient passing parameter sets. Simulating 10,000 values for k_68_ we found its plausible range to be 0.0025 – 0.0087 min^-1^, which ranges from 2-fold less to 1.75-fold more than its calibrated value.

This MPSA generated 4,060 physiologically plausible parameter sets, including parameter sets producing MPS responses within the healthy range (Table S1), above it, and below it. We simulated the kinetic model using these parameter sets following a 3.59-gram leucine bolus and calculated MPS over the 4-hour postprandial period. The parameter sets were then classified as “anabolic sensitive” (n = 3,333) or “anabolic resistant” (n = 727) based on the anabolic resistance threshold (see “Defining Anabolic Resistance” in Methods). The Kolmogorov-Smirnov tests revealed significant differences in the distributions of ten parameters (Figure 2, Figures S2-4, Table S4). These ten parameters were largely consistent with those identified in the p-1 sensitivity analysis of the MPS regression model (see Supplementary Results and Table S2). Two exceptions were splanchnic extraction, which was not varied in the regression analysis, and hepatic glucose production, which only marginally met the *q*<0.05 threshold for inclusion. Of the ten parameters, eight were proximal to the MPS reaction, while the remaining two reflected systemic processes: first-pass splanchnic extraction and hepatic glucose production (from the insulin secretion module).

**Figure 2:**
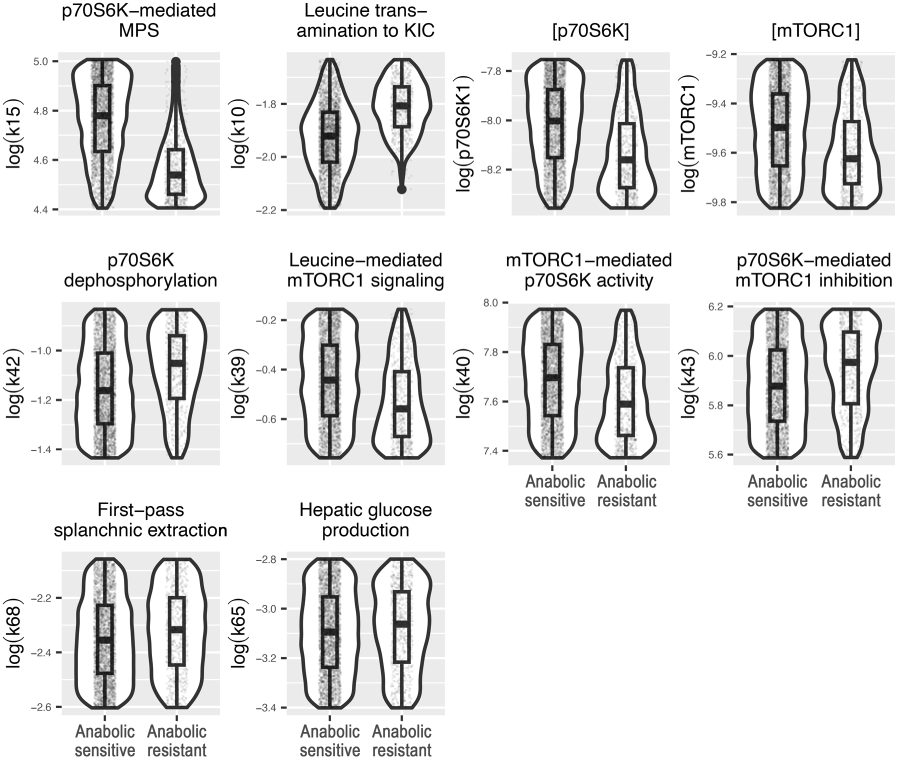
Parameter distributions that significantly differ between the anabolic sensitive and resistant groups. Each panel displays a violin plot showing the parameter distribution density across the range, accompanied by boxplots indicating the median, first and third quartile, and the minimum and maximum values, as well as a scatterplot of the data. The y-axis displays the log10-transformed values of the indicated kinetic model variable.

### Targeted simulations reveal a multifactorial basis of anabolic resistance

We then simulated the model by integrating the putative mechanisms underlying anabolic resistance – individually and in combination, with or without compensatory signaling – using both consensus and worst-case estimates to evaluate their contributions to blunted MPS responses (Table 1). Simulations using the consensus estimates revealed reduced mTORC1 sensitivity (as indicated by p70S6K phosphorylation status) substantially decreased MPS (Figure 3A), closely approximating the MPS response in older adults as calculated from Mitchell et al. (2017) [31] (0.27±0.09 grams of leucine synthesized). Increased first-pass splanchnic extraction and reduced mTOR and p70S6K protein concentrations modestly decreased MPS, while reduced blood flow and impaired insulin signaling had minimal effects (Figure 3A). When all mechanisms were simulated together, the model predicted a substantial decline in MPS, falling below the experimentally observed MPS response in older adults [31]. However, when compensatory signaling was included – simulating the elevated postabsorptive phospho-p70S6K levels observed in older adults – the loss of MPS was recovered to the healthy level. This recovery indicates that reduced mTORC1 sensitivity alone does not account for the age-related loss of MPS when the combined and compensatory effects are considered. Changes in NB closely mirrored those in MPS, as MPB responses remained relatively stable compared to the healthy simulations, however the combined simulation with compensatory signalling retained a slight deficit in NB (Figures 3B and S5).

**Figure 3:**
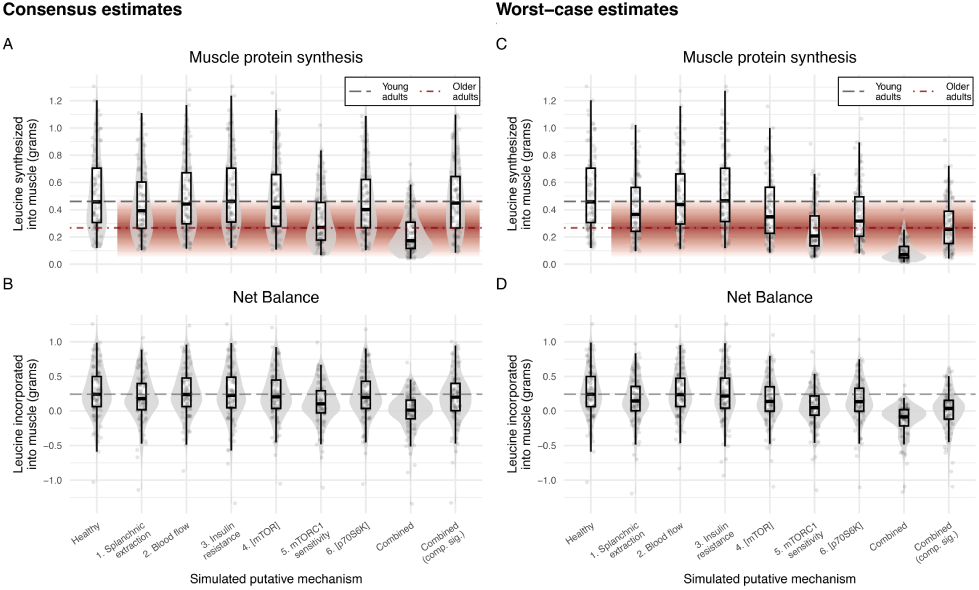
Simulation of putative mechanisms underlying anabolic resistance: Impact on muscle protein synthesis and net balance. Muscle protein synthesis (MPS) and net balance (NB) responses were simulated following a 3.59-gram leucine bolus using either consensus (A, B) or worst-case (C, D) estimates for six putative mechanisms of anabolic resistance (Table 1). Each mechanism was individually simulated by multiplying the relevant model parameter by the value estimated from older adult data. In the “Combined” condition, all six putative mechanisms were simultaneously simulated. The “Combined (comp. sig.)” condition additionally includes compensatory signaling (comp. sig.) to simulate elevated postabsorptive phospho-p70S6K levels. The “Healthy” simulation represents a subset of 100 plausible muscle protein metabolism profiles that met a conservative MPSA criterion for a healthy individual. This same subset was used to simulate each anabolic resistance condition, enabling visualization of the potential distribution of responses across mechanisms. Each simulated condition displays a violin plot showing the distribution density, overlaid with a boxplot (median, first and third quartile, and minimum and maximum values), and individual data points. In the MPS panels, the grey dashed line denotes the MPS response in young adults, as calculated from Mitchell et al. (2015) [30]. The red dot-dashed line and surrounding red gradient represent the mean MPS response and 95% confidence interval for older adults, as calculated from Mitchell et al. (2017) [31]. In the NB panels, the grey dashed line indicates the median NB value of the healthy simulation.

Simulations using the worst-case estimates of the anabolic resistance mechanisms revealed that reduced mTORC1 sensitivity drastically reduced MPS, falling well below the MPS response calculated from Mitchell et al. (2017) [31] (Figure 3C). Increased first-pass splanchnic extraction and reduced mTOR and p70S6K levels markedly reduced MPS, approaching the calculated MPS response [31] (Figure 3C). In contrast, reduced blood flow and impaired insulin-mediated signaling had little effect on MPS. When all mechanisms were simulated together under worst-case conditions, MPS declined substantially falling below the MPS response calculated in older adults [31]. Compensatory signaling restored much of this loss, yielding a MPS response that closely matched levels observed in older adults [31]. As in the consensus simulations, MPB responses were largely unaffected, such that changes in NB closely reflected the changes in MPS (Figures 3D and S5).

Next, we simulated the effect of elevated MPB rates on muscle protein metabolism. Although accurately estimating MPB in older adults is challenging and age-related increases in MPB have not been consistently observed in healthy aging [12], factors such as chronic low-grade inflammation, which contributes to increased MPB [29], and reduced insulin-mediated suppression of MPB [27,28] have been observed in older adults. To evaluate the potential effects of increased MPB, we simulated the model with a 50% increase in the kinetic parameters controlling Akt- and mTORC1-mediated MPB, both independently and in combination with all anabolic resistance mechanisms, with or without compensatory signaling. Increased MPB led to a slight rise in MPS due to greater availability of free intramuscular leucine, but this effect was offset by a larger increase in MPB, resulting in a net reduction in protein balance (Figure S6). When these elevated MPB effects were added to the combined anabolic resistance simulations – with or without compensatory signaling – MPS increased slightly, but the larger rise in MPB further reduced NB.

### Therapeutic strategies to overcome the mechanisms of anabolic resistance

Next, we simulated therapeutic interventions by restoring each dysregulated mechanism to its healthy value under three scenarios: 1) when the mechanism itself was individually dysregulated, 2) when the same magnitude of recovery was applied to other individually dysregulated mechanisms, even though these parameters were not themselves impaired, to evaluate potential compensatory effects, and 3) when all mechanisms were simultaneously dysregulated. The effectiveness of each intervention was evaluated against its ability to recover MPS and NB to the healthy responses.

Under the *consensus estimates*, recovering mTORC1 sensitivity fully restored MPS and NB when applied to individually simulated mechanisms, while recovery of p70S6K levels also substantially restored both outcomes except for dysregulated mTORC1 sensitivity (Figure 4A-C). However, when all anabolic resistance mechanisms were simulated together without compensatory signaling, no single therapeutic target was sufficient to restore MPS or NB. But when we simulated the combined recovery of splanchnic extraction, mTORC1 sensitivity, and both mTOR and p70S6K levels, near-complete recovery of MPS and NB was achieved.

**Figure 4:**
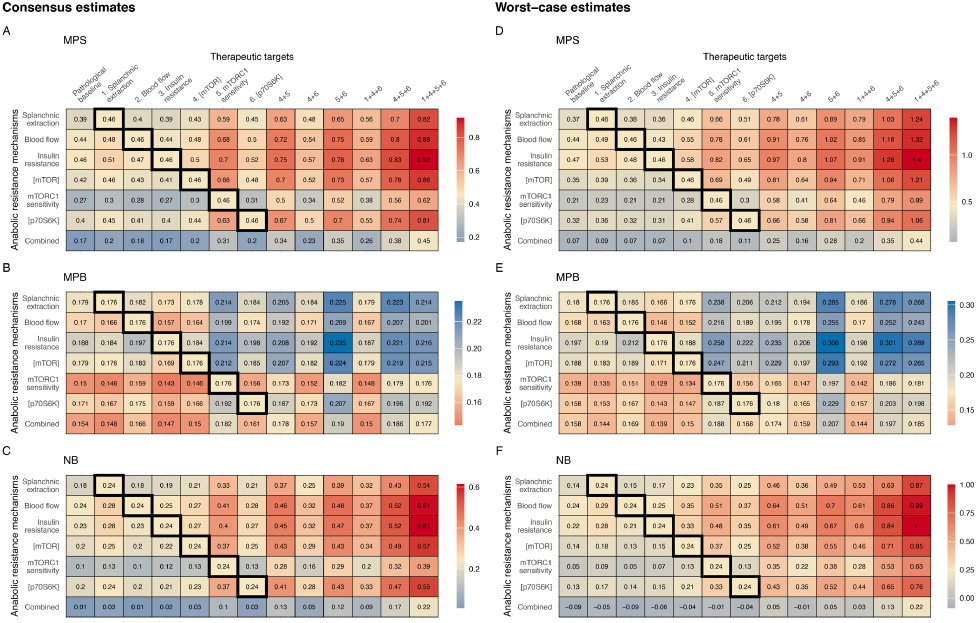
Simulated therapeutic strategies to overcome anabolic resistance and restore muscle protein metabolism. Each therapeutic target was simulated across a subset of 100 model parameter sets by restoring the relevant parameter to its healthy value (i.e., multiplying by the inverse of the pathological estimate from Table 1). Bolded entries in therapeutic targets 1 to 6 indicate cases where restoring a single mechanism, together with the simulation of the respective anabolic resistance mechanism, returned the muscle metabolism response to the healthy level. Combined therapeutic strategies (denoted by “+”) reflect the co-simulation of multiple therapeutic targets. All individual and combined therapeutic strategies were simulated against each anabolic resistance mechanism individually and against all six mechanisms combined. The “Pathological baseline” column shows the median muscle metabolism response across the 100 parameter sets for each anabolic resistance condition without intervention (Figure 3). A heatmap displays the simulated median responses for muscle protein synthesis, muscle protein breakdown, and net balance (NB) across conditions, using either consensus (A-C) or worst-case (D-F) estimates. Gold indicates recovery to the healthy value; blue indicates a response that would cause a reduction in NB compared to the healthy value; red indicates a response that cause produce an increase in NB compared to the healthy value. The numerical value in each cell represents the median muscle metabolism response – expressed in units of grams of leucine synthesized (MPS), degraded (MPB), or net incorporated into muscle (NB) – calculated across the subset of 100 simulated model parameter sets.

When therapeutic strategies were simulated using the *worst-case estimates* (i.e., the measured values deviating most from the healthy state), recovery of mTORC1 sensitivity fully restored MPS and NB when anabolic resistance mechanisms were simulated individually, while recovery of either mTOR or p70S6K levels also largely restored both responses, except for reduced mTORC1 sensitivity (Figure 4D-F). As with the consensus simulations, no single therapeutic target could overcome the combined effects of all anabolic resistance mechanisms. However, combined recovery of splanchnic extraction, mTORC1 sensitivity, and mTOR and p70S6K levels produced a near-complete restoration of MPS and NB.

We then simulated the therapeutic strategies in the presence of compensatory signaling. This analysis was based on the hypothesis that compensatory signaling – reflected by elevated postabsorptive phospho-p70S6K levels in older adults [10,34] – may offset the effects of reduced mTOR and p70S6K concentrations and decreased mTORC1 sensitivity. Accordingly, we simulated these three mechanisms together (“[mTOR] + [p70S6K] + mTORC1 sensitivity”), as well as all anabolic resistance mechanisms together, using the worst-case estimates, in the presence of compensatory signaling and the applied therapeutic strategies (Figure 5). Compensatory signaling alone restored a large portion of MPS and partially restored NB toward healthy levels in both simulated conditions, but neither fully reached the healthy level (healthy values: MPS = 0.46 g leucine synthesized, NB = 0.24 g leucine integrated into muscle; Figure 5). When the three mechanisms were simulated together, several therapeutic strategies in the presence of compensatory signaling successfully restored MPS and NB: recovery of p70S6K levels fully restored the MPS and NB response, and alternative combined strategies including p70S6K recovery could further restore MPS and NB. When all six mechanisms were simulated with compensatory signaling, additional interventions were required to recover both MPS and NB. Two combinations fully recovered both outcomes: 1) recovery of mTOR and p70S6K levels together with mTORC1 sensitivity, and 2) recovery of splanchnic extraction along with mTOR and p70S6K levels. Adding both recovered splanchnic extraction and mTORC1 sensitivity in addition to mTOR and p70S6K recovery produced an even greater increase in MPS and NB.

**Figure 5.**
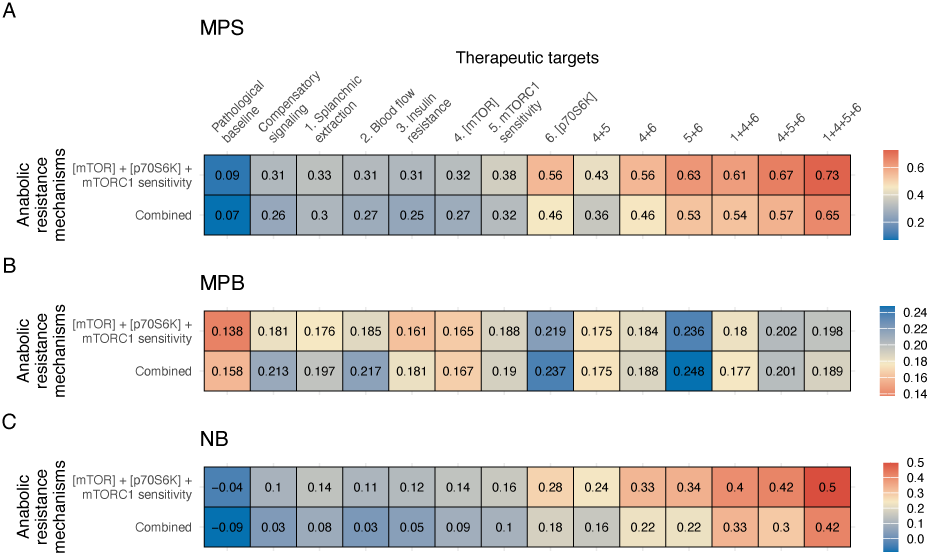
Simulated therapeutic strategies to overcome anabolic resistance and restore protein metabolism in muscle cells with compensatory signaling. The worst-case estimates for reduced mTOR concentration, reduced p70S6K concentration, and reduced mTORC1 sensitivity were simulated simultaneously (“[mTOR] + [p70S6K] + mTORC1 sensitivity”), along with the combined simulation of all six anabolic resistance mechanisms (“Combined”). These simulations were performed without compensatory signaling (“Pathological baseline”), with compensatory signaling to simulate elevated postabsorptive phospho-p70S6K levels, and with compensatory signaling plus individual or combined therapeutic strategies (combinations denoted by “+”). A heatmap displays the simulated median responses for muscle protein synthesis, muscle protein breakdown, and net balance (NB) across conditions. Gold indicates recovery to the healthy value; blue indicates a response that would produce a reduction in NB compared to the healthy value; red indicates a response that would produce an increase in NB compared to the healthy value. The numerical value in each cell represents the median muscle metabolism response – expressed in units of grams of leucine synthesized (MPS), degraded (MPB), or net incorporated into muscle (NB) – calculated across the subset of 100 simulated model parameter sets. For reference, the healthy values for MPS, MPB, and NB were 0.47 g synthesized, 0.154 g degraded, and 0.27 g net incorporated, respectively.

## Discussion

In this study we used a computational modeling approach to systematically evaluate the mechanisms that most likely contribute to anabolic resistance. We first conducted a global sensitivity analysis of a previously developed kinetic model of leucine-mediated signaling and muscle protein metabolism [10] to identify mechanisms that most strongly control MPS, MPB, and NB. The analysis revealed that MPS and MPB are primarily controlled by their proximal regulatory mechanisms, whereas NB is largely influenced by MPS regulation. Next, we performed an exploratory analysis simulating an EAA feeding intervention to evaluate mechanisms contributing to anabolic resistance in older adults [31], classifying simulated responses as either anabolic sensitive or resistant. Several parameters and signaling protein concentrations differed significantly between groups, particularly those controlling MPS. We then conducted a targeted analysis to evaluate the extent to which putative mechanisms underlying anabolic resistance could explain the reduced MPS observed with aging. Model simulations showed that, when considered individually, reduced mTORC1 sensitivity alone could account for anabolic resistance. However, elevated post-absorptive phospho-p70S6K levels – a phenomenon we term “compensatory signaling” – are observed in older adults, and model simulations indicated that this response fully restored MPS when using the consensus estimates, even in the presence of all other putative mechanisms. Under the worst-case estimates, compensatory signaling was insufficient to recover MPS, and when all putative mechanisms were considered together, the model closely replicated the experimentally observed anabolic resistance MPS response. These simulations demonstrate that the putative mechanisms operate together to suppress MPS, while compensatory signaling plays a critical role in partially or fully restoring anabolic capacity depending on the severity of underlying impairments. Finally, we simulated therapeutic interventions aimed at restoring muscle metabolism in anabolic resistance. While single-target approaches (e.g., increasing mTORC1 sensitivity, enhancing mTOR or p70S6K levels) were generally effective when only individual mechanisms were impaired, they proved insufficient when multiple dysregulated processes were simultaneously present. In contrast, restoring MPS when all dysregulated processes were simultaneously present required a multifactorial therapeutic strategy targeting several pathways. These findings highlight the complex, multifactorial nature of age-related anabolic resistance and motivate novel hypotheses about the most likely dysregulated mechanisms, providing a foundation for future experimental validation and multi-targeted therapeutic development.

### Main findings and implications

This study examined strategies to restore the feeding-mediated MPS response in older adults, with a focus on *individual feedings* rather than adjustments to feeding variables such as dose or scheduling. In our simulations, a single 15-g EAA feeding (containing 3.59 g leucine) was modelled. While increasing the simulated dose or changing the scheduling of feeding could be considered as alternatives to improving the MPS responses in older adults, evidence from human studies suggests these strategies provide limited benefits: in older adults, a 10-g EAA feeding maximizes myofibrillar MPS rates with no further increases at 20 g or 40 g EAA [18]. Similarly, studies report that more frequent, smaller feedings (4 × 3.75 g EAA every 45 minutes) [41] or a leucine top-up after an initial EAA bolus (15 g EAA + 3 g leucine at 90 minutes) [31] do not meaningfully enhance MPS rates in older adults compared to a single 15 g EAA bolus. Together, these findings suggest that overcoming anabolic resistance in older adults requires targeting mechanisms that control the response to individual feedings, rather than modifying nutritional feeding strategies alone.

In contrast, some studies indicate that increasing protein dose relative to body weight [14] or enriching the EAA dose with leucine [42] can restore MPS in older adults to levels observed in younger adults. While these compensatory strategies may offset the blunted anabolic response, they do not explain *why* higher protein or leucine doses are necessary. The underlying factors driving this reduced sensitivity likely contribute to muscle loss, such that directly addressing them offers a more sustainable solution for overcoming anabolic resistance. Accordingly, this study focused on identifying the mechanisms responsible for the diminished MPS response to individual feedings and proposes therapeutic strategies to restore this response.

A well-documented mechanism of anabolic resistance is the diminished sensitivity of mTORC1 signaling to protein or amino acid ingestion [17,18,26]. The findings from our naïve approach support this notion because we observed widespread dysregulation in leucine-mediated signaling, including leucine-mediated mTORC1 activity, mTORC1-mediated p70S6K phosphorylation, p70S6K dephosphorylation, and p70S6K-mediated negative feedback on mTORC1. The model simulations from the targeted analysis further highlight the critical contribution of reduced mTOR1 sensitivity to losses in MPS. While experimental studies have yet to pinpoint the specific mechanisms responsible for reduced mTORC1 signaling sensitivity in anabolic resistance [42], our results suggest that a broad dysregulation of the mTORC1 signaling machinery may underlie this phenomenon.

One factor that contributes to reduced mTORC1 signaling sensitivity is physical activity. A synergistic relationship exists between nutrient- and exercise-mediated mTORC1 signaling and MPS, in which exercise enhances muscle sensitivity to protein feeding [43] and improves insulin-mediated amino acid delivery to muscle, thereby supporting mTORC1 signaling [44]. Conversely, physical inactivity desensitizes muscle to feeding-mediated MPS responses [45]. While our simulations focused solely on the feeding response, reduced levels of physical activity in older adults may contribute to the diminished mTORC1 signaling sensitivity seen in anabolic resistance [46–48]. This combination contributes to the progressive desensitization of mTORC1 signaling to anabolic stimuli, exacerbating anabolic resistance.

Our simulations from the naïve approach revealed that mTORC1 and p70S6K concentrations were significantly reduced in the anabolic-resistant group, suggesting a potential mechanism for the blunted MPS response in aging. Consistent with these findings, previous research shows that older adults have 50% lower mTOR and p70S6K levels compared to younger adults that was concomitant with a diminished EAA feeding-induced myofibrillar FSR [18]. Despite these reductions in protein content, older adults exhibit increased postabsorptive phospho-p70S6K^T389^_,_ a potential compensatory mechanism. This increased postabsorptive phospho-p70S6K is seen both in experimental studies [17,18,27,49] and in our previous modeling simulations [10]. This compensation likely occurs to partially offset the reduced mTORC1 and p70S6K concentrations. Indeed, our targeted analysis found that compensatory increases in postabsorptive phospho-p70S6K can largely restore both MPS and NB responses. However, this compensatory signaling was sufficient to restore muscle metabolism only in the consensus estimate simulations. Even under the consensus estimate simulations, considerable variability remained in the predicted MPS response – ranging both above and below the healthy level – indicating that a less effective compensatory response could lead to anabolic resistance. This persistent postabsorptive phosphorylation reduces the dynamic range for further postprandial increases [17,18]. This impaired signaling response may contribute to the diminished MPS observed in older adults following amino acid ingestion.

The compensations in muscle metabolism induced by elevated p70S6K phosphorylation in aging could be offset by two negative feedback mechanisms. First, phosphorylated p70S6K inhibits insulin signaling via IRS1 phosphorylation, reducing insulin sensitivity [50–52]. Second, phospho-p70S6K phosphorylates mTORC1 at Ser2448, further suppressing mTORC1 activation [53–56]. These feedback pathways likely act together to decrease the sensitivity of p70S6K to anabolic stimuli and may in part lead to anabolic resistance. Aged rats similarly exhibit elevated postabsorptive phosphorylation at both p70S6K and its downstream target, ribosomal protein S6 (rpS6), further supporting this mechanism [57]. Interestingly, inhibiting mTORC1 signaling with rapamycin in aged rats reduced phospho-p70S6K and phospho-rpS6 levels, resulting in improved muscle mass maintenance and reduced age-related muscle loss [57]. Given that rapalogs inhibiting mTOR activity have been proposed to improve muscle health with age [58], our modeling suggests this could further restore skeletal muscle sensitivity to anabolic stimuli. However, careful titration of doses is needed, particularly in older adults with co-morbidities and/or low physical activity, to avoid excessive suppression of mTOR signaling and anabolism [59]. Nevertheless, our results suggest an alternative intervention to inhibiting mTORC1 – which we hypothesize primarily targets the compensatory response to reduced p70S6K levels – could be increasing absolute mTORC1 and/or p70S6K. Our model suggests that increasing mTOR or p70S6K levels, individually, were two of the most efficacious therapeutic approaches in recovering muscle metabolism. In particular, therapeutic targeting of p70S6K levels was highly effective in restoring MPS and NB in the presence of compensatory signaling. This approach of targeting mTOR or p70S6K protein levels could restore the capacity for phosphorylation increases, reduce postabsorptive mTORC1 signaling, and ultimately mitigate anabolic resistance.

While this study has identified key potential therapeutic targets to overcome anabolic resistance, a critical next step is to determine which of these is *druggable*, i.e., can be modified by pharmacological agents (cf. [37]). Pharmacotherapies for sarcopenia have a long history of targeting cellular signaling pathways by binding to intracellular or membrane receptors [38]. These therapies primarily focus on modulating signaling pathway activity to either increase MPS or decrease MPB [38]. However, our results suggest that restoring mTORC1 sensitivity and recovering mTOR and p70S6K protein levels may represent more efficacious strategies to overcome anabolic resistance – strategies that differ fundamentally from conventional approaches targeting anabolic or catabolic signalling pathways in muscle. Pharmacological agents such as MHY1485 [60] and rapamycin [57] can enhance or repress mTOR activity, respectively, offering opportunities to modulate mTORC1 sensitivity. Pharmacological strategies aimed at restoring signaling protein levels in older adult muscle represent a novel and promising avenue.

Determining effective pharmacological strategies to modulate mTOR or p70S6K protein levels can be informed by bioinformatics-based approaches (cf. [61]). Specifically, these proteins are encoded by the *MTOR* and *RPS6KB1* genes, such that plausible strategies include identifying factors that upregulate their expression, such as transcription factors (TFs) and cofactors, and targeting these to increase activity. Conversely, repressors of gene expression, such as microRNAs (miRNA) and long non-coding RNAs (lncRNAs), could be inhibited to relieve suppression. For example, miR-199a-3p is a miRNA known to repress *MTOR* expression [62] and could be inhibited to enhance *MTOR* expression [63]. One issue with targeting miRNAs is that multiple *MTOR*-specific miRNAs have been identified, such that targeting just one might be insufficient. Regarding *RPS6KB1*, estrogen receptor alpha (*ESR1*) is one of the few TFs studied for its role in controlling *RPS6KB1* gene expression [64,65], making it a potential target to increase *RPS6KB1* mRNA levels [61]. Additionally, miR-200b [66] and miR-223-3p [67] represent candidate miRNAs whose inhibition may upregulate *RPS6KB1* expression. While these targets for modulating mTOR and p70S6K protein levels are speculative, pharmacological approaches that inhibit TF-DNA interactions, modulate enhancer activity, or repress miRNAs have been developed and successfully applied in other biological contexts [68–70]. It is important to note that these interventions primarily affect mRNA levels; therefore, proteomic analyses are essential to confirm corresponding changes at the protein level and assess the functional impact of these strategies.

Our study features the following noteworthy limitations. First, we evaluated simulated responses and their approximation to anabolic resistance against two studies [30,31]. Given the substantial heterogeneity in anabolic resistance and dysregulated muscle protein metabolism with aging [14,71,72], future studies should further validate and refine our model using larger, individual-level datasets. Additionally, future human clinical trials could benefit from measuring the critical contributors to anabolic resistance identified here to generate robust datasets. Second, our model was developed using data from males. Some [73], but not all [74], studies suggest that aging-related dysregulation of muscle protein metabolism may exhibit sexual dimorphism. Therefore, future studies should include females to confirm the generalizability of our findings. Finally, our results focus on muscle protein metabolism, which most likely affects muscle mass. While losses of muscle mass, driven by dysregulations in protein turnover, often precedes declines in muscle function [75], the relationship between mass and function is not always direct [74].

## Conclusions

We report a novel modeling approach to efficiently investigate key variables associated with age-related anabolic resistance. The initial sensitivity analysis revealed widespread dysregulation of reactions controlling mTORC1 activity in the anabolic-resistant group, along with reduced levels of mTORC1 and p70S6K. Simulating the putative mechanisms underlying anabolic resistance suggested that these mechanisms do not act in isolation; rather, they act synergistically to suppress MPS. Based on these results, we simulated therapeutic interventions to restore muscle metabolism. Targeting individual mechanisms proximal to mTORC1 signalling (e.g., mTORC1 sensitivity, mTOR or p70S6K levels,) effectively restored MPS and NB when only individual impairments were simulated. However, these single-target strategies failed to fully restore MPS when all mechanisms were impaired simultaneously. Under this more realistic condition – reflecting what we suggest is likely occurring in older adults – a multifactorial therapeutic approach was required to recover anabolic responses to feeding. These findings motivate new hypotheses regarding the combinatorial mechanisms underlying anabolic resistance and offer insight into future experimental studies and therapeutic strategies to mitigate feeding-mediated anabolic resistance. Furthermore, our framework could also be applied to other models of muscle loss characterized by similar dysregulations, such as inactivity-induced anabolic resistance [45], or distinct mechanisms, such as increased MPB in cachexia [76].

## Acknowledgements

We thank Drs. Robert Wolfe and Arny Ferrando for their thoughtful discussion in improving this work. This work was supported by a Natural Sciences and Engineering Research Council of Canada (NSERC) Collaborative Research and Training Experience scholarship to T.J.M. and a NSERC Discovery Grant to D.C.C. (RGPIN 06004-2014). The funders had no role in study design, data collection and analysis, decision to publish, or preparation of the manuscript. DDC is currently supported by a National Institutes of Health (NIH) Clinical Research Loan Repayment Award. The content is solely the responsibility of the authors and does not necessarily represent the official views of the National Institutes of Health. The findings are presented clearly, honestly, and without fabrication, falsification, or inappropriate data manipulation.

