## Supplementary material for "Exploring the multifactorial causes and therapeutic strategies for anabolic resistance in sarcopenia: A systems modeling study"

### Supplementary Information

#### Supplementary Methods

##### *Kinetic model of leucine-mediated signaling and muscle protein metabolism*

The McColl and Clarke kinetic model simulates skeletal muscle protein metabolism in response to leucine ingestion in healthy adults and contains four interconnected modules: 1) the mTOR signaling module, 2) the leucine kinetic module, 3) the digestive system module, and 4) the insulin secretion module [1]. The mTOR signaling module represents the dynamic behavior of key signaling proteins within the mTOR network in response to insulin and leucine stimulation. Given leucine's central role as an activator of mTORC1, a dedicated leucine kinetics module was incorporated to simulate its physiological dynamics. This module captures leucine absorption into muscle cells and its subsequent metabolism, including incorporation into muscle protein, transamination to $\alpha$-ketoisocaproic acid (KIC), and KIC oxidation. The digestive system module accounts for leucine intake and absorption into the bloodstream. It incorporates factors such as gastric emptying and absorption kinetics when leucine is consumed in a low-caloric, low-volume solution in the postprandial state. Additionally, it includes the ileal digestibility of leucine and its first-pass splanchnic extraction. The insulin secretion module simulates the ultradian secretion of insulin from the pancreas, governed by regulatory feedback loops between glucose and insulin. A comprehensive description of the model can be found in McColl and Clarke [1].

The kinetic model consists of four compartments – stomach, gut, blood plasma, and skeletal muscle – and includes 34 molecular species, comprising 11 proteins, 13 post-translationally modified proteins, and 10 metabolite pools [1]. It incorporates 63 kinetic parameters, of which 60 are adjustable and three are constrained, along with three delay parameters controlling insulin and glucose production [1]. The kinetic model was constructed using nonlinear ordinary differential equations to simulate the rate of change of each molecular species in units of moles [1].

##### *Multi-parameter sensitivity analysis*

We applied the multi-parameter sensitivity analysis (MPSA) following the approach by Zi [2]. The sensitivity analysis was conducted in four key steps:

1. Sampling – A total of 100,000 parameter sets were generated using Latin hypercube sampling (MATLAB function: *lhsdesign*) to ensure comprehensive coverage within

predefined parameter uncertainty bounds. Each parameter was sampled on a $\log_{10}$ scale, ranging from two-fold above to two-fold below its calibrated value from McColl and Clarke [1]. In the MPSA of the kinetic model (see corresponding Results and Supplementary Results section), five parameters controlling leucine transport and insulin dynamics (k1, k2, k3, k4, k68) were held constant.

2. Simulation – The kinetic model was simulated following a 3.5-gram leucine bolus for each sampled parameter sets.
3. Filtering – Model outputs were evaluated against conservative experimental ranges following leucine ingestion to identify *physiologically plausible* parameter sets in healthy adults (Figure S1). While the McColl and Clarke [1] model used a single parameter set to represent a healthy individual, substantial inter-individual variability in physiological and biochemical factors suggests that model parameters likely vary across a healthy population. For example, differences in muscle mass, composition, and distribution [3], insulin sensitivity [4], amino acid absorption kinetics [5], and intracellular responsiveness to anabolic stimuli [6] have been documented in healthy adults. To account for this variability, we applied conservative acceptability criteria for key experimentally measured outputs to generate a virtual population of healthy adults. Parameter sets that met all filtering criteria (detailed below) were deemed physiologically plausible for a healthy adult ("plausible"), while those that failed at least one criterion were considered physiologically implausible ("implausible"). Although many of these "implausible" parameter sets were numerically feasible and did not cause simulation error, their outputs fell outside the defined physiological bounds for a healthy adult.
4. Analysis – The subset of plausible parameter sets was further examined using multiple regression and the Kolmogorov-Smirnov (KS) test to determine key contributors to model behavior.

After identifying plausible parameter sets, we re-simulated the kinetic model using each parameter set to evaluate key muscle protein metabolism outcomes, including muscle protein synthesis (MPS, the amount of leucine incorporated into skeletal muscle protein), muscle protein breakdown (MPB, the amount of leucine degraded from skeletal muscle protein), and net balance (NB, the difference between MPS and MPB). MPS was measured as the area under the curve (AUC) of the MPS reaction ($r_{15}$), MPB was measured as the sum of the AUC of the Akt- ($r_{69}$) and mTORC1-mediated

MPB ($r_9$) reactions, and NB was calculated as the difference between MPS and MPB (NB = MPS – MPB).

*MPSA: qualitative criteria for parameter set filtering*

To identify plausible parameter sets, we applied conservative qualitative criteria based on the calibration data and model simulations in McColl and Clarke [1]. These criteria featured four variables that reflected the dynamic responses of model species to leucine ingestion. The four variables are defined as follows:

1. Postabsorptive concentration of model species at the end of the burn-in period: $[species]_{t=0}$
2. Timing of peak concentration of the model species: $t_{\Delta[species]_{peak}}$
3. Peak concentration difference of the model species: $\Delta[species]_{peak}$
4. Return to postabsorptive concentration of the model species: $[species]_{postabsorptive}$

The passing boundaries for criterion 3 (peak concentration difference, $\Delta[species]_{peak}$) were calculated as a percent difference relative to the postabsorptive value ($[species]_{t=0}$) and the peak concentration ($[species]_{peak}$) for each simulation, as described in Equations 1 and 2:

$$\Delta[species]_{peak_{upper}} = \left([species]_{peak} - [species]_{t=0}\right) \times (1 + \delta) + [species]_{t=0} \quad (1)$$

$$\Delta[species]_{peak_{lower}} = \left([species]_{peak} - [species]_{t=0}\right) \times (1 - \delta) + [species]_{t=0} \quad (2)$$

Where Equation 1 defines the upper bound, Equation 2 defines the lower, and $\delta$ represents the fractional percent change (e.g., $\delta$ = 0.5 for a ±50% change).

For phospho-protein species, the passing criteria for criteria 1 and 3 were adjusted. The $[species]_{t=0}$ criterion was calculated as the percent of protein phosphorylated relative to the total protein concentration, as shown in Equation 3:

$$[species]_{t=0} = \frac{[protein_{i_{phos}}]}{[protein_{i_{total}}]} \times 100\% \quad (3)$$

where $[protein_{i_{phos}}]$ is the concentration of phosphorylated $protein_i$, and $[protein_{i_{total}}]$ is the total concentration of $protein_i$. The $\Delta[species]_{peak}$ criterion was calculated as the phospho-protein fold-change between the peak phospho-protein concentration and the postabsorptive phospho-protein concentration, as shown in Equation 4:

$$\Delta[species]_{peak} = \frac{[protein_{i_{phos}}]}{[protein_{i_{phos_{t=0}}}]} \quad (4)$$

where $[protein_{i_{phos_{t=0}}}]$ is the postabsorptive concentration of phosphorylated $protein_i$.

For the MPS model species, we defined the following separate criterion in the MPSA, $FSR_{simulated}$, which was computed as the AUC of the MPS reaction, simulating the MPS response over the postprandial period.

Three model species were evaluated against all three MPSA criteria, three model species were evaluated against the first three criteria, and MPS was evaluated against criterion 2 and $FSR_{simulated}$ (Figure S1, Table S1).

*MPSA: multiple regression*

We applied multiple regression analysis to the plausible parameter sets to evaluate the relationship between model parameters and muscle protein metabolism outcomes. Regression models were generated using the MATLAB function *fitlm* for the following response variables:

1. MPS ~ parameter values
2. MPB ~ parameter values
3. NB ~ parameter values

Both raw and $\log_{10}$-transformed response variables were analyzed for MPS and MPB, whereas only raw values were analysed for NB because it is calculated as the difference between MPS and MPB. For each response variable, regression models were developed using either $\log_{10}$-transformed or z-score normalized parameter values. This resulted in four models for MPS and MPB, but only two models for NB:

1. Raw outcome ~ $\log_{10}$(parameter values)
2. Raw outcome ~ z-score(parameter values)
3. $\log_{10}$(outcome) ~ $\log_{10}$(parameter values)
4. $\log_{10}$(outcome) ~ z-score(parameter values)

Each regression model was first run with only main effect terms. Interaction terms were then added in increments of 5 (e.g., 5, 10, 15, etc.) to determine the most predictive model. The interaction terms were selected iteratively based on the most significant covariates (i.e., kinetic model parameters) from the main effect-only model. Model goodness-of-fits were assessed using adjusted $R^2$ and Akaike information criterion (AIC). A model was considered significantly more predictive if its AIC (Equation 6) was reduced by at least 10 compared to the previous best-fit model [7].

$$\Delta AIC = AIC_i - AIC_{min} \tag{6}$$

where $AIC_i$ is the AIC value of the currently evaluated model and $AIC_{min}$ is the lowest AIC value of the current subset of model.

*MPSA: p-1 sensitivity analysis*

Once the optimal regression model was determined for each muscle protein metabolism outcome, we performed a p-1 sensitivity analysis to evaluate the parameters with greatest influence on model responses. Following the approach of Clarke et al. [8], we systematically removed from the full regression model each model parameter one at a time, including both its main effect and any associated interaction terms. Model fit was evaluated using adjusted $R^2$ and AIC. Parameters with the largest ΔAIC had the most influence on model response, whereas parameters with a ΔAIC less than 10 were considered to have a negligible effect.

*MPSA: Kolmogorov-Smirnov test*

The two-sample Kolmogorov-Smirnov (KS) test was used to determine whether parameter sets originated from the same distribution. We performed the test using the R function *ks.test*, which calculated the KS statistic and the associated p-value. To account for multiple comparisons, we applied false discovery rate (FDR) correction using the Benjamini-Hochberg method (R function: *p.adjust*) to compute the *q*-value. A *q*-value less than 0.05 indicated a statistically significant difference between parameter distributions.

#### Supplementary Results

##### *Multi-parameter sensitivity analysis of the kinetic model*

We varied all but five kinetic model parameters within a two-fold range. The five fixed parameters included those controlling leucine transport through the digestive tract ($k_1$), leucine excretion ($k_2$), leucine absorption ($k_3$), leucine-mediated insulin secretion ($k_4$), and first-pass splanchnic extraction ($k_{68}$), which were kept at their calibrated values. Parameters $k_{1\text{-}3}$ and $k_{68}$ define the model's input function (i.e., leucine-mediated stimulation), such that varying them produced excessive implausible parameter sets. Similarly, varying $k_4$ caused large deviations in plasma insulin dynamics, further increasing the number of implausible parameters sets. Fixing these parameters ensured that sufficient plausible parameter sets could be generated for analysis.

This initial MPSA produced 1,155 physiologically plausible parameter sets for healthy adults. Among the passing parameter distributions, we found that the parameters controlling plasma leucine absorption into muscle ($k_6$) and intracellular leucine transamination to α-KIC ($k_{10}$) exhibited distributions for the passing parameter values that did not span the full two-fold range:

- $k_6$: calibrated value = 0.0331 min$^{-1}$; two-fold range = 0.0166 – 0.0662 min$^{-1}$; passing range = 0.0264 – 0.0457 min$^{-1}$
- $k_{10}$: calibrated value = 0.0117; two-fold range = 0.0058 – 0.0233 = passing range, 0.0069 – 0.0230

These restricted distributions indicate that the model performance is particularly sensitive to $k_6$ and $k_{10}$ relative to the others. Therefore, we performed the MPSA again but with reduced ranges for $k_6$ (0.75- to 1.42-fold of the calibrated value) and $k_{10}$ (0.55- to 2-fold of the calibrated value). This refined MPSA yielded 2,663 physiologically plausible parameter sets for healthy adults. We used the 2,663 plausible parameter sets for subsequent analysis.

##### *Multiple regression to determine model parameters most contributing to muscle protein metabolism*

Using the plausible parameter sets from the MPSA, we developed multiple regression models to determine key parameters influencing muscle metabolism outcomes (MPS, MPB, and NB). We first simulated the kinetic model using the plausible parameter sets to generate MPS, MPB, and NB outcome measures over a 3-hour period following the ingestion of a 3.5-gram leucine bolus. The simulated outcomes varied as follows:

- MPS: 0.180 – 0.540 grams of leucine incorporated into muscle (included as a MPSA criterion)
- MPB: 0.009 – 1.234 grams of leucine degraded from muscle
- NB: -0.982 to 0.520 grams of leucine (net muscle loss to gain)

Among the four regression models developed for MPS and MPB, we found the $\log_{10}$-transformed outcome measures with $\log_{10}$-transformed parameter values were the most predictive (Table S2). Among the two regression models developed for NB, we found the raw outcomes with $\log_{10}$-transformed parameter values were the most predictive (Table S2).

The best-performing regression models for MPS, MPB, and NB include the top 35, 45, and 30 parameters from the main-effect-only models as interaction terms, respectively (see Table S3 for a representative analysis of the sequential addition of interaction terms to determine the most predictive model).

The p-1 sensitivity analysis was conducted using the most predictive models (Table 2). For MPS and MPB, the analysis revealed that parameters most proximal to their respective reactions had the greatest influence on their outcomes. For NB, the most influential parameters were a combination of those affecting MPS and MPB, but parameters more proximal to MPS had a greater influence on the NB response.

We then performed the p-1 sensitivity analysis using the best-performing regression models from the other three versions for each of the protein metabolism outcome measures (data not shown). The top 10 most influential model parameters were identical for MPS, and only slightly varied in ordering for MPB and NB. This consistency across models suggests that the findings are robust, regardless of the regression model version used.

### Supplementary Figures

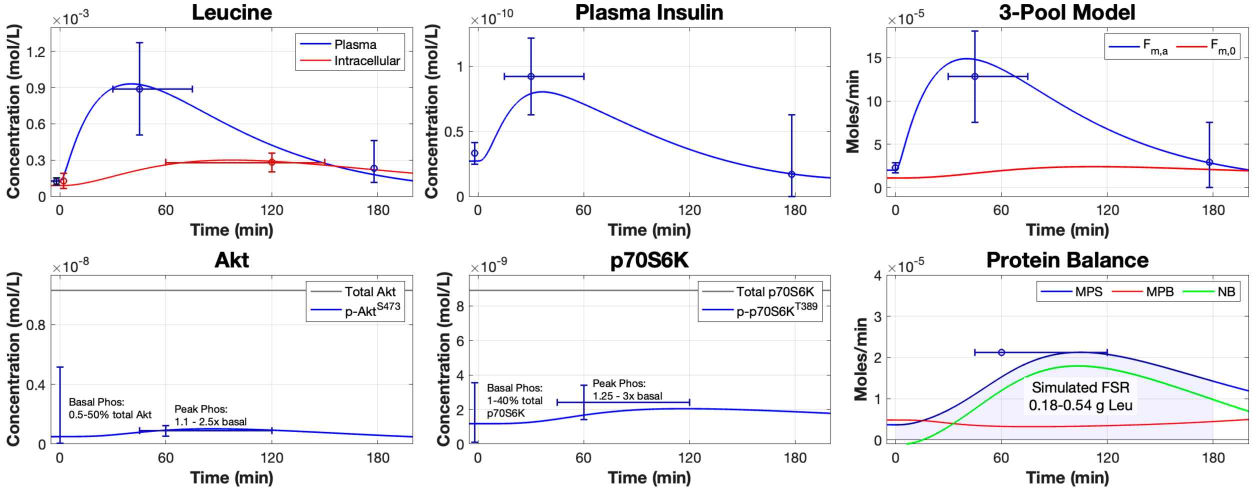


**Figure S1:** MPSA filtering criteria. Conservative ranges of experimental data following leucine ingestion used to classify parameter sets in the MPSA as physiologically "plausible" or "implausible" for a healthy adult. The selection criteria for the model dynamic of plasma leucine, intracellular leucine, plasma insulin, $F_{m,a}$, phospho-Akt, phospho-p70S6K, and MPS were coded according to four components: 1) postabsorptive levels, 2) peak concentration, 3) timing of peak concentration, and 4) return to postabsorptive levels. The selection criteria for MPS was also coded for a conservative range of total MPS via the area under the curve following leucine ingestion (i.e., simulated FSR). Each of these model species were coded for a combination of these components. The error bars in each plot indicate the conservative data range for each component. Simulated parameter sets from the MPSA that passed through all conservative ranges were classified as "plausible" parameter sets.

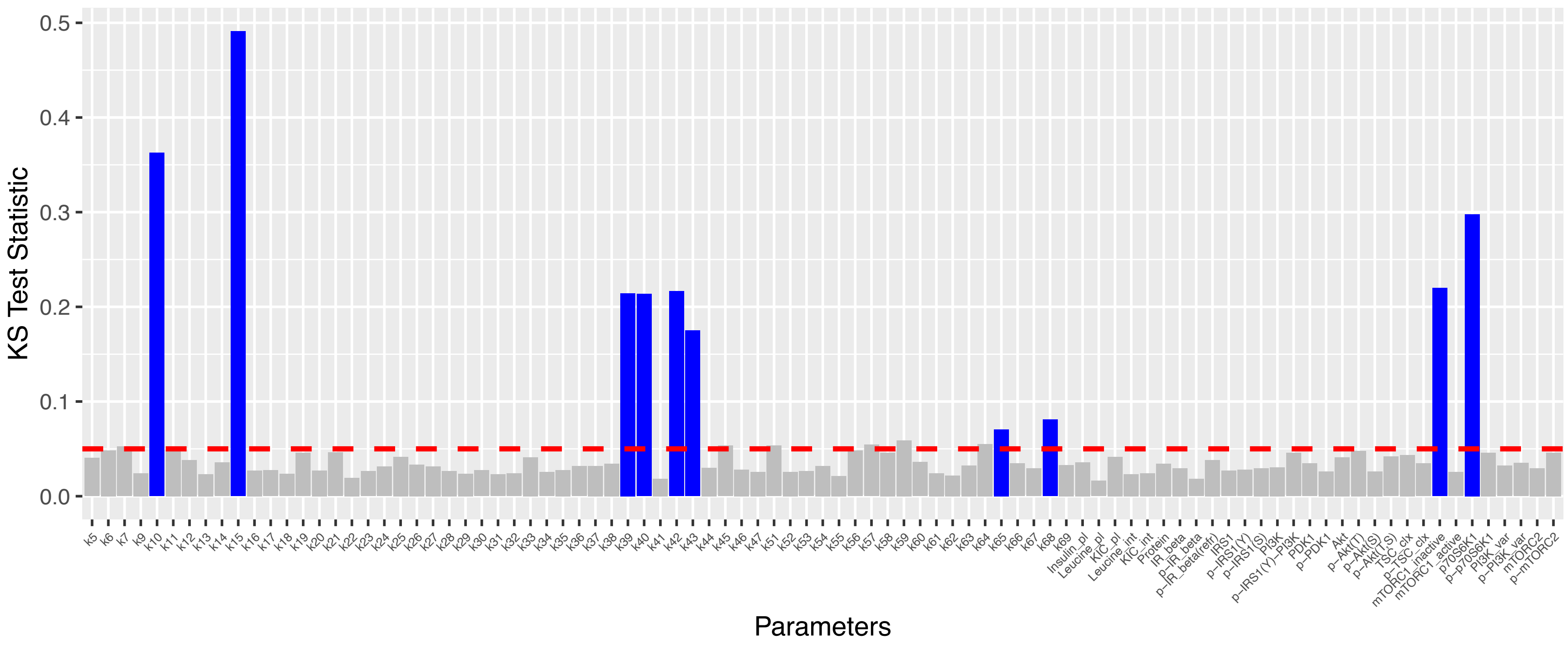


**Figure S2:** Two-sample KS test comparing parameter distributions between the anabolic sensitive and resistance groups. The red dashed line represents the statistical significance threshold after false discovery rate correction ($q<0.05$). Parameter distributions that differ significantly between the anabolic sensitive and resistant groups are shown in blue, while non-significant distributions are shown in grey. See Table S4 for statistical outputs.

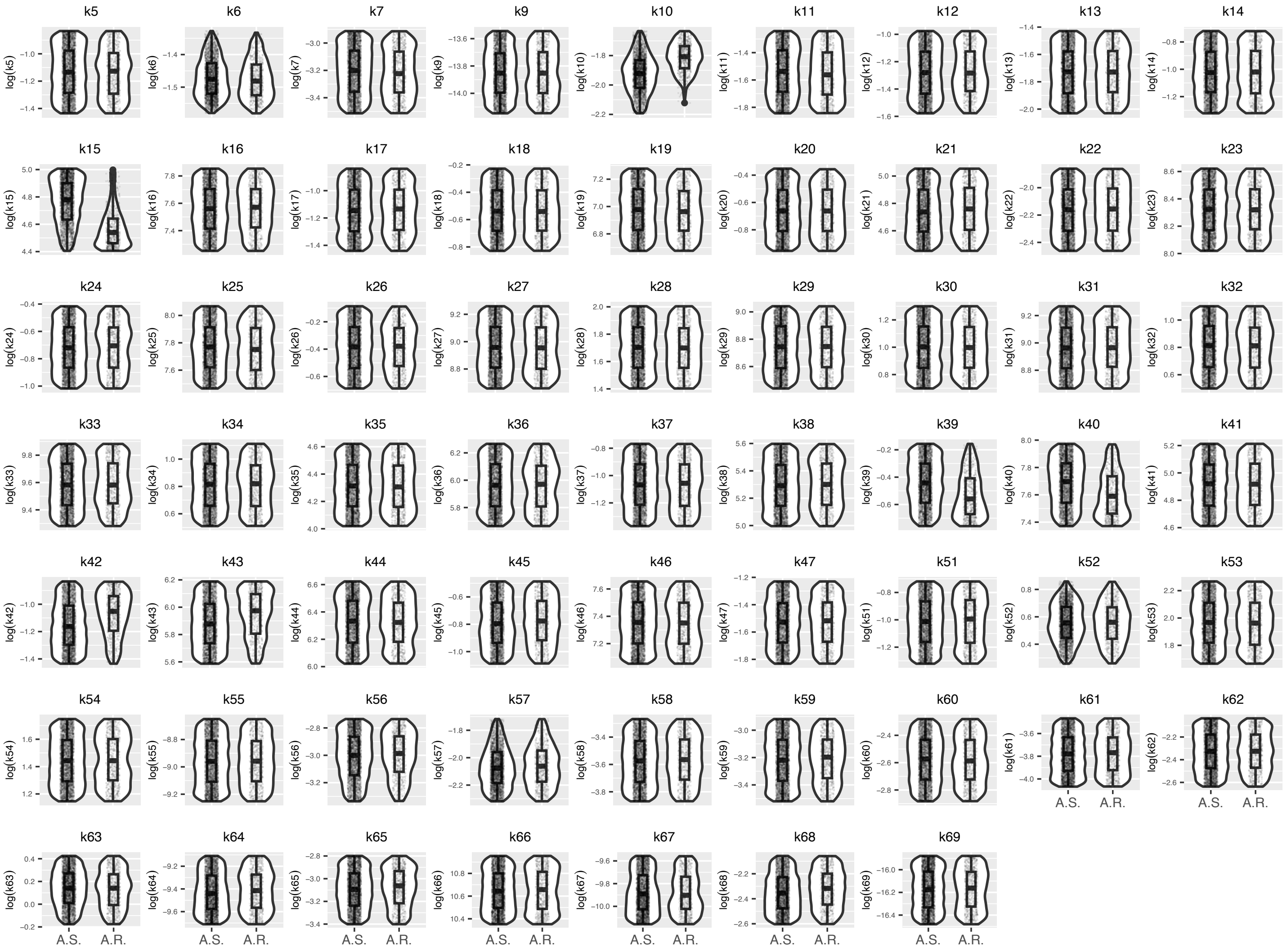


**Figure S3:** Kinetic parameter distributions between the anabolic sensitive and resitant groups. Each panel displays a violin plot showing the parameter distribution density across the range, accompanied by boxplots indicating the median, first and third quartile, and the minimum and maximum values, as well as a scatterplot of the data. The y-axis label corresponds to the specific kinetic model variable, log10 transformed. A.R., anabolic resistant group; A.S., anabolic sensitive group.

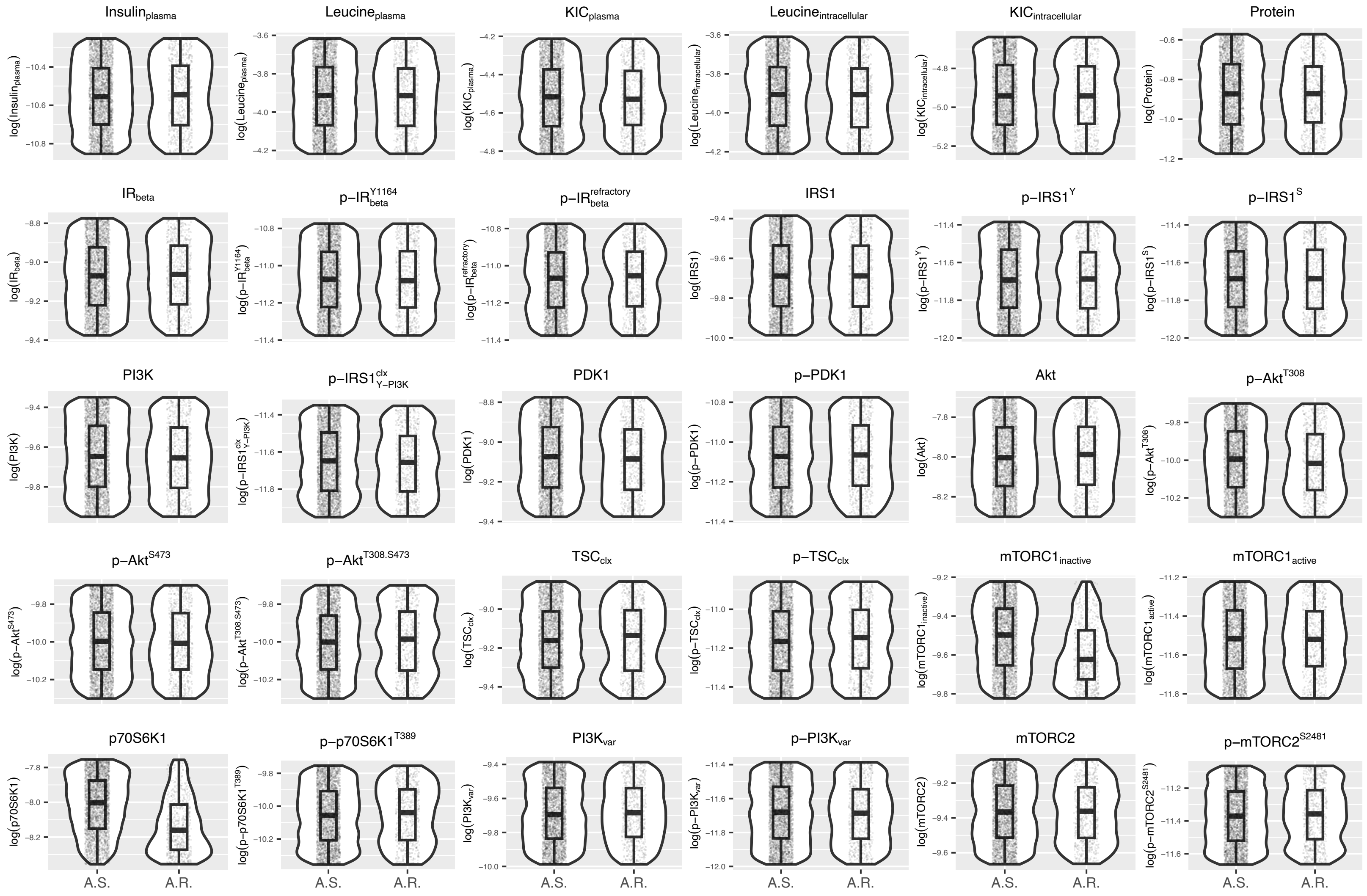


**Figure S4:** Initial concentration parameter distributions between the anabolic sensitive and resistant groups. Each panel displays a violin plot showing the parameter distribution density across the range, accompanied by boxplots indicating the median, first and third quartile, and the minimum and maximum values, as well as a scatterplot of the data. The y-axis label corresponds to the specific kinetic model variable, log10 transformed. A.R., anabolic resistant group; A.S., anabolic sensitive group.

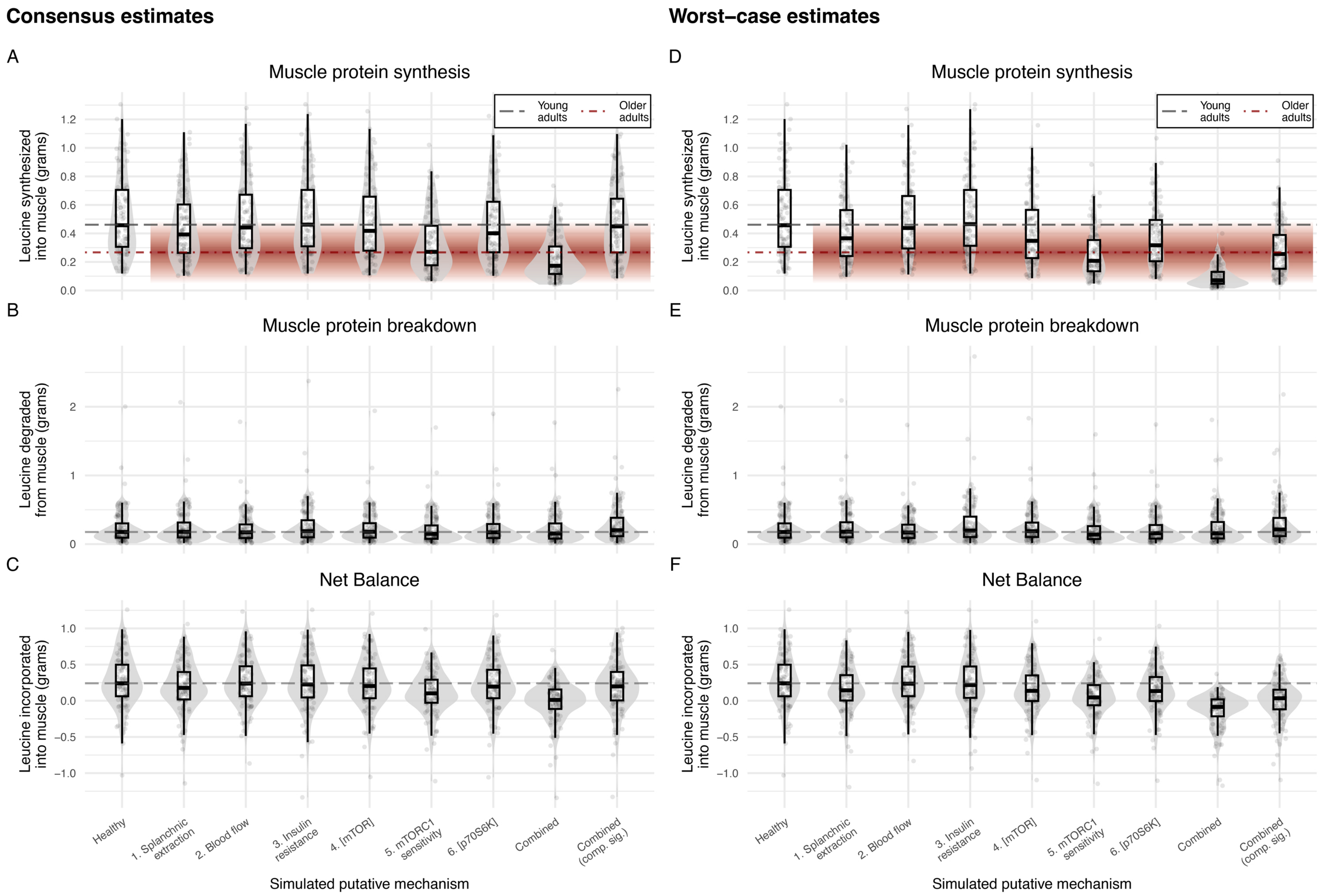

Consensus estimates
Worst–case estimates
A
Muscle protein synthesis
D
B
Muscle protein breakdown
E
C
Net Balance
F
Young adults
Older adults
Leucine synthesized into muscle (grams)
Leucine degraded from muscle (grams)
Leucine incorporated into muscle (grams)
Healthy
1. Splanchnic extraction
2. Blood flow
3. Insulin resistance
4. [mTOR]
5. mTORC1 sensitivity
6. [p70S6K]
Combined
Combined (comp. sig.)
Simulated putative mechanism


216

**Figure S5:** Simulation of the putative mechanisms underlying anabolic resistance: Impact on muscle protein synthesis, muscle protein breakdown, and net balance. Muscle protein synthesis (MPS), muscle protein breakdown (MPB), and net balance (NB) responses were simulated following a 3.59-gram leucine bolus using either consensus (A, B, C) or worst-case (D, E, F) estimates of the putative mechanisms of anabolic resistance, as reported in Table 1. Each of the six mechanisms were individually simulated by multiplying the relevant model parameter by the value estimated from older adult data. In the "Combined" conditions, all six putative mechanisms were simulated together, with compensatory signaling (comp. sig.) included in the "Combined (comp. sig.) condition to simulate elevated levels of postabsorptive phospho-p70S6K.

The "healthy" simulation represents a subset of 100 plausible muscle protein metabolism responses that passed a conservative MPSA criterion for a healthy individual. This same subset was used to generate each anabolic resistance simulation, illustrating the potential distribution of responses across mechanisms.

Each simulated condition displays a violin plot showing the parameter distribution density, overlaid with a boxplot indicating the median, first and third quartile, and the minimum and maximum values, as well as a scatterplot of the data.

In the MPS panels, the grey horizontal dashed line represents the MPS response in young adults, as calculated from Mitchell et al. (2015)[9]. The red horizontal dot-dashed line, together with the surrounding red gradient, represents the mean MPS response and its 95% confidence interval in older adults, as calculated from Mitchell et al. (2017)[10]. In the MPB and NB panels, the gray dashed line represents the median value of the healthy simulation.

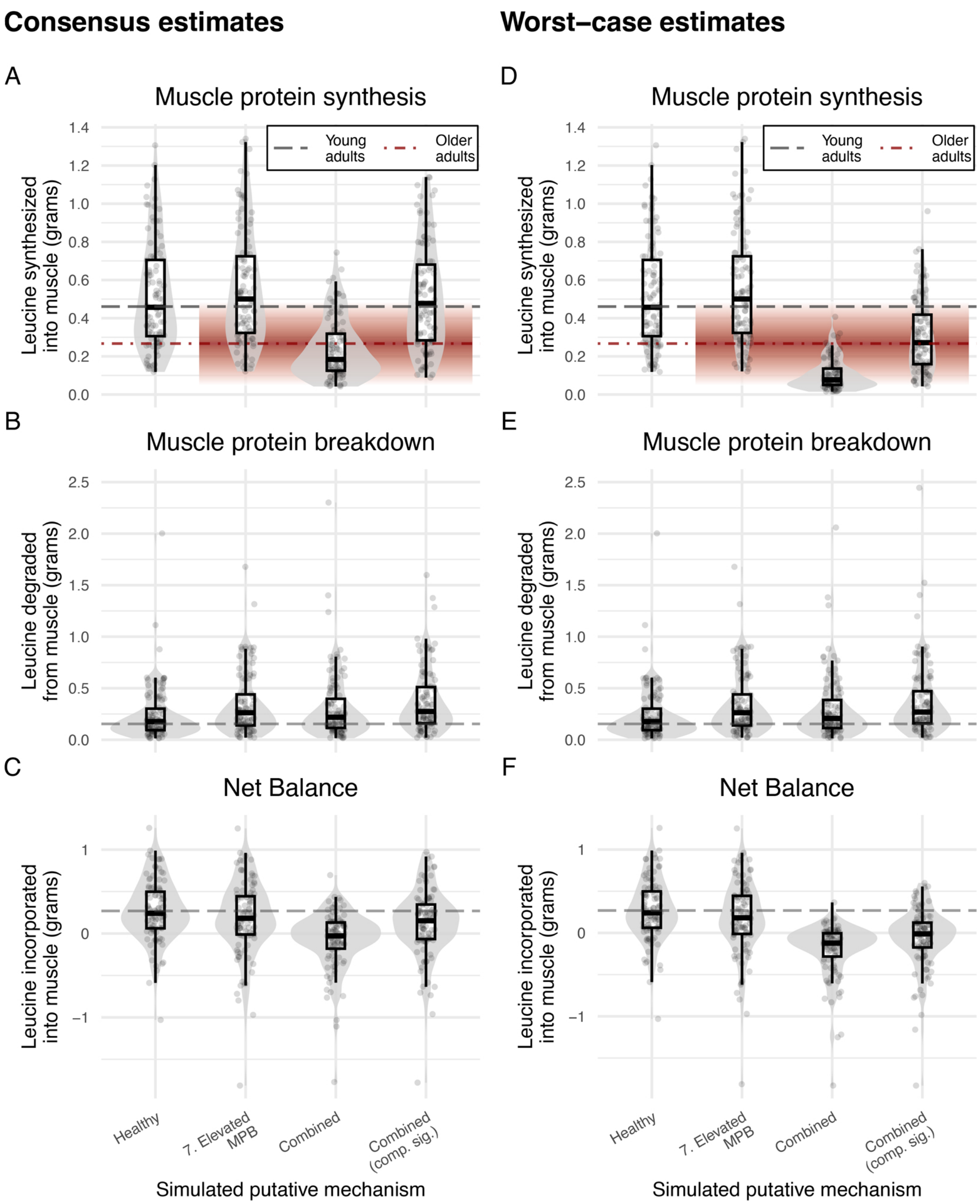
Consensus estimates
Worst–case estimates
A
Muscle protein synthesis
D
Muscle protein synthesis
Young adults
Older adults
Leucine synthesized into muscle (grams)
B
Muscle protein breakdown
E
Muscle protein breakdown
Leucine degraded from muscle (grams)
C
Net Balance
F
Net Balance
Leucine incorporated into muscle (grams)
Healthy
7. Elevated MPB
Combined
Combined (comp. sig.)
Simulated putative mechanism

**Figure S6:** Simulation of elevated muscle protein breakdown rates together with the mechanisms underlying anabolic resistance: Impact on muscle protein synthesis, muscle protein breakdown, and net balance. Muscle protein synthesis (MPS), muscle protein breakdown (MPB), and net balance (NB) responses were simulated following a 3.59-gram leucine bolus with a 50% elevation to the model parameters controlling Akt- and mTORC1-mediated MPB together with either consensus (A, B, C) or worst-case (D, E, F) estimates of the putative mechanisms of anabolic resistance, as reported in Table 1. Elevated MPB rates were simulated individually, together with the combination of all six putative mechanisms of anabolic resistance ("Combined"), or together all six putative mechanisms and the addition of compensatory signaling (comp. sig.) to simulate elevated levels of positabsorptive phospho-p70S6K ("Combined (comp. sig.)").

The "healthy" simulation represents a subset of 100 plausible muscle protein metabolism responses that passed a conservative MPSA criterion for a healthy individual. This same subset was used to generate each anabolic resistance simulation, illustrating the potential distribution of responses across mechanisms.

Each simulated condition displays a violin plot showing the parameter distribution density, overlaid with a boxplot indicating the median, first and third quartile, and the minimum and maximum values, as well as a scatterplot of the data.

In the MPS panels, the grey horizontal dashed line represents the MPS response in young adults, as calculated from Mitchell et al. (2015)**[9]**. The red horizontal dot-dashed line, together with the surrounding red gradient, represents the mean MPS response and its 95% confidence interval in older adults, as calculated from Mitchell et al. (2017)**[10]**. In the MPB and NB panels, the gray dashed line represents the median value of the healthy simulation.

### Supplementary Tables

**Table S1:** MPSA filtering criteria.

| Model species | MPSA criterion | Exp. data | Simulated data [1] | MPSA passing range | |
|---|---|---|---|---|---|
| | | | | Allowed deviance | Calculated range |
| Plasma leucine | $[species]_{t=0}$ | 123 μM[346] | 132 μM | ±25% | 92–154 μM |
| | $t_{\Delta[species]_{peak}}$ | 45 min[346] | 42 min | -15, +30 min | 30–75 min |
| | $\Delta[species]_{peak}$ | 890 μM[346] | 960 μM | ±50% | 507–1,273 μM |
| | $[species]_{postab.}$ | 231 μM[346] | 189 μM | ±200% | 116–462 μM |
| Intracellular leucine | $[species]_{t=0}$ | 128 μM[347] | 86 μM | ±50% | 64–192 μM |
| | $t_{\Delta[species]_{peak}}$ | 120 min[347] | 98 min | -60, +30 min | 60–150 min |
| | $\Delta[species]_{peak}$ | 280 μM[347] | 290 μM | ±50% | 204–356 μM |
| Plasma insulin | $[species]_{t=0}$ | 33 pM[346] | 26 pM | ±25% | 25–41 pM |
| | $t_{\Delta[species]_{peak}}$ | 30 min[346] | 36 min | -15, +30 min | 15–60 min |
| | $\Delta[species]_{peak}$ | 92 pM[346] | 75 pM | ±50% | 63–122 pM |
| | $[species]_{postab.}$ | N/a | 17 pM | Conserv$ | <63 pM |
| $F_{m,a}$ | $[species]_{t=0}$ | 23 μmol/min[346] | 20 μmol/min | ±25% | 17–29 μmol/min |
| | $t_{\Delta[species]_{peak}}$ | 45 min[346] | 42 min | -15, +30 min | 30–75 min |
| | $\Delta[species]_{peak}$ | 130 μmol/min[346] | 146 μmol/min | ±50% | 83–210 μmol/min |
| | $[species]_{postabs.}$ | 29 μmol/min[346] | 29 μmol/min | Conserv$ | <75 μmol/min |
| p-Akt$^{S\wedge}$ | $[species]_{t=0}$# | N/a | 26.5 % phos. | | 0.5–50% phos. |
| | $t_{\Delta[species]_{peak}}$ | 60 min[346] | 91 min | -15, +60 min | 45–120 min |
| | $\Delta[species]_{peak}$* | 1.41 F.C.[346] | 1.46 F.C. | | 1.1–2.5 F.C. |
| p-p70S6K | $[species]_{t=0}$# | N/a | 13.3% phos. | | 1–40% phos. |
| | $t_{\Delta[species]_{peak}}$ | 90 min[346] | 116 min | -45, +60 min | 45–150 min |
| | $\Delta[species]_{peak}$* | 1.85 F.C.[346] | 1.71 F.C. | | 1.25–3 F.C. |
| MPS | $FSR_{simulated}$ | 0.36 g leu[346] | 0.36 g leu | ±50% | 0.18–0.54 g leu |
| | $t_{\Delta[species]_{peak}}$ | 60 min[346] | 105 min | -15, +60 min | 45–120 min |

$Conservative estimates were used for the return of species to postabsorptive concentrations. These were calculated as half the difference between $[species]_{peak}$ and $[species]_{t=0}$:

$$Conserv.estimate = \left([species]_{peak} - [species]_{t=0}\right) \times 0.5 + [species]_{t=0}$$

^p-Akt$^{S}$ represents the sum of both serine phosphorylated Akt species: p-Akt$^{S473}$ and p-Akt$^{S473,T308}$

#The $[species]_{t=0}$ for phospho-protein species was calculated as the percent of protein phosphorylated relative to total protein concentration.

*The $\Delta[species]_{peak}$ for phospho-protein species was calculated as the fold change between the peak phospho-protein concentration and the postabsorptive phospho-protein concentration.

$[species]_{postabs.}$, $[species]_{postabsorptive}$; F.C., fold-change; phos., phosphorylated.

**Table S2:** P-1 sensitivity analysis results using the most predictive multiple regression models for MPS, MPB, and NB. Each of the MPS, MPB, and NB models includes interactions terms for the top 35, 45, and 35 parameters from the main effect-only models, respectively. The top 10 parameters from the analyses are presented. ΔAIC is the difference between the p-1 regression model and the most predictive model (i.e., all terms included).

| Parameter removed | Parameter description | Adjusted $R^2$ | AIC | ΔAIC |
|---|---|---|---|---|
| | *$log_{10}$(MPS) ~ $log_{10}$(parameters)* | | | |
| All terms included | | 0.987 | -14,131 | |
| Param. removed: | | | | |
| k15 | Muscle protein synthesis | 0.327 | -3726 | 10405 |
| p70S6K1 | p70S6K initial concentration | 0.663 | -5565 | 8566 |
| k10 | Leucine transamination to α-KIC | 0.720 | -6060 | 8071 |
| k40 | mTORC1-mediated p70S6K phos. | 0.756 | -6428 | 7703 |
| k42 | p70S6K dephosphorylation | 0.768 | -6565 | 7566 |
| $mTORC1_{inactive}$ | mTORC1 initial concentration | 0.782 | -6725 | 7406 |
| k39 | Leucine-mediated mTORC1 activity | 0.794 | -6880 | 7251 |
| k43 | p70S6K-mediated mTORC1 feedback | 0.848 | -7688 | 6443 |
| k11 | α-KIC reamination to leucine | 0.936 | -9996 | 4135 |
| k14 | α-KIC oxidation | 0.941 | -10222 | 3909 |
| | *$log_{10}$(MPB) ~ $log_{10}$(parameters)* | | | |
| All terms included | | 0.991 | -9,477 | |
| Param. removed: | | | | |
| Protein | Protein initial concentration | 0.798 | -1225 | 8252 |
| k9 | p-$Akt^{T308}$-mediated blunting of MPB | 0.868 | -2367 | 7110 |
| Akt | Akt initial concentration | 0.874 | -2490 | 6987 |
| k28 | p-$Akt^{T308}$ dephosphorylation | 0.883 | -2683 | 6794 |
| PDK1 | PDK1 initial concentration | 0.892 | -2883 | 6594 |
| k27 | PDK1-mediated $Akt^{T308}$ phosphorylation | 0.904 | -3195 | 6282 |
| k26 | p-PDK1 dephosphorylation | 0.917 | -3582 | 5895 |
| k25 | p-$IRS1^{Y}$-PI3K-mediated PDK1 phos. | 0.920 | -3692 | 5784 |
| PI3K | PI3K initial concentration | 0.942 | -4579 | 4898 |
| IRS1 | IRS1 initial concentration | 0.944 | -4668 | 4808 |
| | *NB ~ $log_{10}$(parameters)* | | | |
| All terms included | | 0.955 | -9,399 | |
| Param. removed: | | | | |
| k15 | Muscle protein synthesis | 0.726 | -4595 | 4804 |
| k10 | Leucine transamination to α-KIC | 0.844 | -6090 | 3309 |
| p70S6K1 | p70S6K initial concentration | 0.865 | -6476 | 2923 |
| $mTORC1_{inactive}$ | mTORC1 initial concentration | 0.875 | -6681 | 2718 |
| k39 | Leucine-mediated mTORC1 activity | 0.880 | -6781 | 2618 |
| k40 | mTORC1-mediated p70S6K phos. | 0.894 | -7128 | 2271 |
| Protein | Protein initial concentration | 0.896 | -7160 | 2239 |
| k42 | p70S6K dephosphorylation | 0.896 | -7166 | 2232 |
| k43 | p70S6K-mediated mTORC1 feedback | 0.899 | -7263 | 2136 |
| k9 | p-$Akt^{T308}$-mediated blunting of MPB | 0.906 | -7447 | 1951 |

Param., parameter; Phos., phosphorylation.

**Table S3:** This table provides a representative example of the iterative process used to determine the most predictive regression model for protein metabolism outcomes. The $\log_{10}$(MPS) ~ $\log_{10}$(parameters) regression model was initially fit with main effect terms only, followed by the sequential addition of interaction terms for the most significant parameters. The number (N) of regression model parameters and goodness-of-fit measures, including adjusted $R^2$ and Akaike Information Criterion (AIC), are reported. The model incorporating main effect terms and interactions of the top 35 parameters exhibited the lowest AIC, indicating the best predictive performance. A similar approach was used for MPB and NB (data not shown).

| **Regression model: $\log_{10}$(MPS) ~ $\log_{10}$(parameters)** | **N model parameters** | **Adjusted $R^2$** | **AIC** |
|---|---|---|---|
| Main effect terms only | 90 | 0.957 | -11,528 |
| Main effect terms + interactions of top 5 parameters | 100 | 0.961 | -11,777 |
| Main effect terms + interactions of top 10 parameters | 135 | 0.963 | -11,888 |
| Main effect terms + interactions of top 15 parameters | 195 | 0.967 | -12,095 |
| Main effect terms + interactions of top 20 parameters | 280 | 0.973 | -12,575 |
| Main effect terms + interactions of top 25 parameters | 390 | 0.980 | -13,227 |
| Main effect terms + interactions of top 30 parameters | 525 | 0.986 | -14,114 |
| *Main effect terms + interactions of top 35 parameters* | *685* | *0.987* | *-14,131* |
| Main effect terms + interactions of top 40 parameters | 870 | 0.987 | -14,049 |
| Main effect terms + interactions of top 45 parameters | 1080 | 0.987 | -14,015 |

**Table S4.** Model parameters with significant differences between the anabolic sensitive and resistance groups. Model parameters with a q-value < 0.05, as determined by the 2-sample Kolmogorov-Smirnov (KS) test, are shown.

| **Model parameter** | **KS statistic** | ***p*-value** | ***q*-value** |
|---|---|---|---|
| p70S6K-mediated MPS | 0.491 | $9.70\times10^{-127}$ | $9.31\times10^{-125}$ |
| Leucine transamination to $\alpha$-KIC | 0.362 | $5.43\times10^{-69}$ | $2.64\times10^{-67}$ |
| p70S6K initial concentration | 0.298 | $6.54\times10^{-47}$ | $2.09\times10^{-45}$ |
| mTORC1 initial concentration | 0.221 | $6.33\times10^{-26}$ | $1.52\times10^{-24}$ |
| Phospho-p70S6K dephosphorylation | 0.217 | $3.44\times10^{-25}$ | $6.61\times10^{-24}$ |
| Leucine-mediated mTORC1 signaling | 0.214 | $2.03\times10^{-24}$ | $3.25\times10^{-23}$ |
| mTORC1-mediated p70S6K activity | 0.213 | $3.34\times10^{-24}$ | $4.59\times10^{-23}$ |
| p70S6K-mediated mTORC1 inhibition | 0.176 | $1.45\times10^{-16}$ | $1.73\times10^{-15}$ |
| First-pass splanchnic extraction | 0.082 | $5.76\times10^{-4}$ | $6.14\times10^{-3}$ |
| Hepatic glucose production | 0.070 | $5.16\times10^{-3}$ | $4.95\times10^{-2}$ |
